# Integrative multi-omic analysis identifies tumor-intrinsic p38 as a driver of immune exclusion in human epithelial cancers

**DOI:** 10.64898/2026.06.11.731716

**Authors:** Rebekah E. Dadey, Krishna B. Singh, Rose Doerfler, Riley Santiago, Brian Isett, Christopher Deitrick, Carl Kim, Sarah Newman, Marion Joy, Katelyn Smith, Carly Reeder, Armando P. Signore, Ernest M. Meyer, Tullia C. Bruno, Aditi Kulkarni, Qaingqiang Gu, Chen Zhang, Aatur D. Singhi, Raja R. Seethala, Adam C. Soloff, Rajeev Dhupar, Heath D. Skinner, Laura P. Stabile, Lazar Vujanović, Robert L. Ferris, Jason J. Luke, Dan P. Zandberg, Riyue Bao

## Abstract

Patients with tumors not responding to immune-checkpoint inhibition (ICI) often harbor a non-T cell-inflamed tumor microenvironment, characterized by the absence of IFN-γ-associated CD8^+^ T cells and dendritic cell activation. While the role of p38 mitogen-activated protein kinases (MAPKs) in regulating dendritic and myeloid cells is established, the tumor-intrinsic immunomodulatory function of p38 remains underexplored. Here, we identify tumor cell-intrinsic p38 signaling as a target candidate associated with immune exclusion and reduced immunotherapy response. In human papillomavirus-negative head and neck squamous carcinoma (HNSCC), molecular analysis of 395 tumor tissues revealed a p38-centered network enriched in non-T cell-inflamed tumors. Multi-cancer single-cell RNA sequencing analysis of over 200,000 cells further identifies p38 activation as a potential immune-exclusion program across multiple epithelial tumor types, including HNSCC and lung squamous cell carcinoma (LUSC), supported by tissue validation in ∼250 human biospecimens using multispectral imaging and digital spatial profiling. Functional studies demonstrate that p38 knockdown or pharmacologic inhibition in HNSCC and LUSC cell lines increases T cell migration, with CXCL16 identified as a chemokine mediator in vitro; neutralization of CXCL16 attenuated this effect. Together, these findings identify tumor-intrinsic p38 activation as a driver of immune exclusion in epithelial cancers and support its potential as a therapeutic target to overcome immunotherapy resistance.

## INTRODUCTION

Immunotherapy with immune-checkpoint inhibition (ICI) has improved outcomes for cancer; however, most patients do not benefit. From a developmental therapeutics perspective, the biology underpinning lack of treatment response is a critical to identify high priority combination approaches. The T cell-inflamed tumor microenvironment (TME), characterized by CD8^+^ T cells, PD-L1 and type I/II interferon (IFN) gene expression, is an important prognostic and predictive biomarker for ICI response(1). Tumor mutational burden (TMB)(2) also dictates response with some oncogenic pathways known to be strongly associated with immunotherapy resistance. These pathways, or more adequately described as tumor-intrinsic mechanisms of immune exclusion, are causally linked to the non-T cell-inflamed phenotype(3). Tumor-cell intrinsic pathways may impact anti-tumor immunity through multiple mechanisms, but previous studies have emphasized reduced recruitment of antigen processing cells via suppression of tumor-derived chemokine gradients(3). We and others identified WNT/β-catenin signaling as the first tumor-intrinsic mechanism of immune exclusion, initially in melanoma(4) and then across tumors(5). Other notable tumor-intrinsic pathways include MYC, PI3K (PTEN loss), IDH1, and STK11 (6–11). Moreover, we described a pan-cancer molecular atlas for tumor-intrinsic mechanisms of immune exclusion using bulk RNA sequencing (RNAseq), confirming our previous findings surrounding WNT signaling and nominating further potential therapeutic targets(12).

Thousands of clinical trials have been completed investigating combinations of anti-PD1 with various mechanistic agents, however only a handful of combinations have become robust strategies in clinic(13). While preclinical murine models have justified the pursuit of these trials, fewer studies have taken an unbiased discovery approach using human tissues to identify combination treatments. The identification of tumor-cell intrinsic molecular signaling as a driver of non-T cell-inflamed tumors therefore provides a human centric and rational framework to investigate the genomics and transcriptomics of immunotherapy resistance with the potential to identify novel therapeutic targets to overcome ICI resistance.

Here we identify a previously unknown role for tumor cell p38 signaling in suppressing anti-tumor immunity that is amenable to targeted inhibition in clinic. p38 mitogen-activated protein kinases (MAPK) are stress mediators strongly associated with progression of several tumor types, with known roles regulating proliferation and senescence as well as apoptosis(14, 15). Within the immune compartment, p38 is known to cross-talk with c-Jun N-terminal kinase (JNK) signaling(15), induce proinflammatory/immunosuppressive myeloid cells and cytokines, and regulating dendritic cell (DC) maturation, type I IFN responses, and antigen presentation(16–18). Within stromal cells of the TME, p38 has also been identified to suppress adaptive immune responses(19). In this study, we designed a hypothesis-driven, multi-omic discovery and validation framework leveraging independent cohorts across transcriptomics, proteomics, spatial profiling with human biospecimen validation, and in vitro functional experiments (**Fig. S1**). Beginning with an unbiased pathway analysis in human papillomavirus (HPV)-negative head and neck squamous cell carcinoma (HNSCC) as a clinically relevant model, we identified and systematically validated a novel tumor cell-intrinsic role for p38 MAPK, specifically MAPK14, in immune exclusion across multiple epithelial cancer types including lung squamous cell carcinoma (LUSC), lung adenocarcinoma (LUAD), and renal clear cell carcinoma (RCC). These data emphasize the potential for unbiased discovery and validation from human biospecimens to identify novel therapeutic combinations and facilitate drug repurposing strategies to overcome ICI resistance in cancer immunotherapy.

## RESULTS

### Tumor-intrinsic signaling pathways are associated with immune exclusion

To understand molecular correlates of immune exclusion within clinically relevant biomarker-defined populations, we pursued HNSCC as a model where stratification of clinical care by the presence of the HPV dictates treatment. We performed an initial analysis of RNAseq data from 481 tumors from The Cancer Genome Atlas (TCGA) HNSCC cohort(20), in concert with pathway analysis, to identify oncogenic pathways differentially activated in non-T cell-inflamed tumors. Given the etiologic impact of HPV in HNSCC, we focused on the 395 HPV-negative tumors (**Fig. 1A**). Using a T cell-inflamed gene expression signature(5, 12, 21, 22), we observed 36% *versus* 30% of tumors expressing the T cell-inflamed *versus* non-inflamed phenotype, respectively. We detected 67 pathways significantly activated in non-inflamed relative to inflamed tumors (z-score≥1.95, *P*<0.05) from causal network analysis(23) (**Fig. 1B**). The activation of upstream regulators was predicted by Ingenuity Pathway Analysis (IPA)’s causal networks (23). We referred to such upstream regulators as “pathways” throughout the manuscript to distinguish them from single genes or molecules. Functional annotation of the pathways revealed two main themes, with 43 pathways representing the MAPK, HIF-1, and T cell-receptor regulation of apoptosis, and 24 representing IL6 pathway and oxidative stress (**Fig. 1B**).

**Figure 1.**
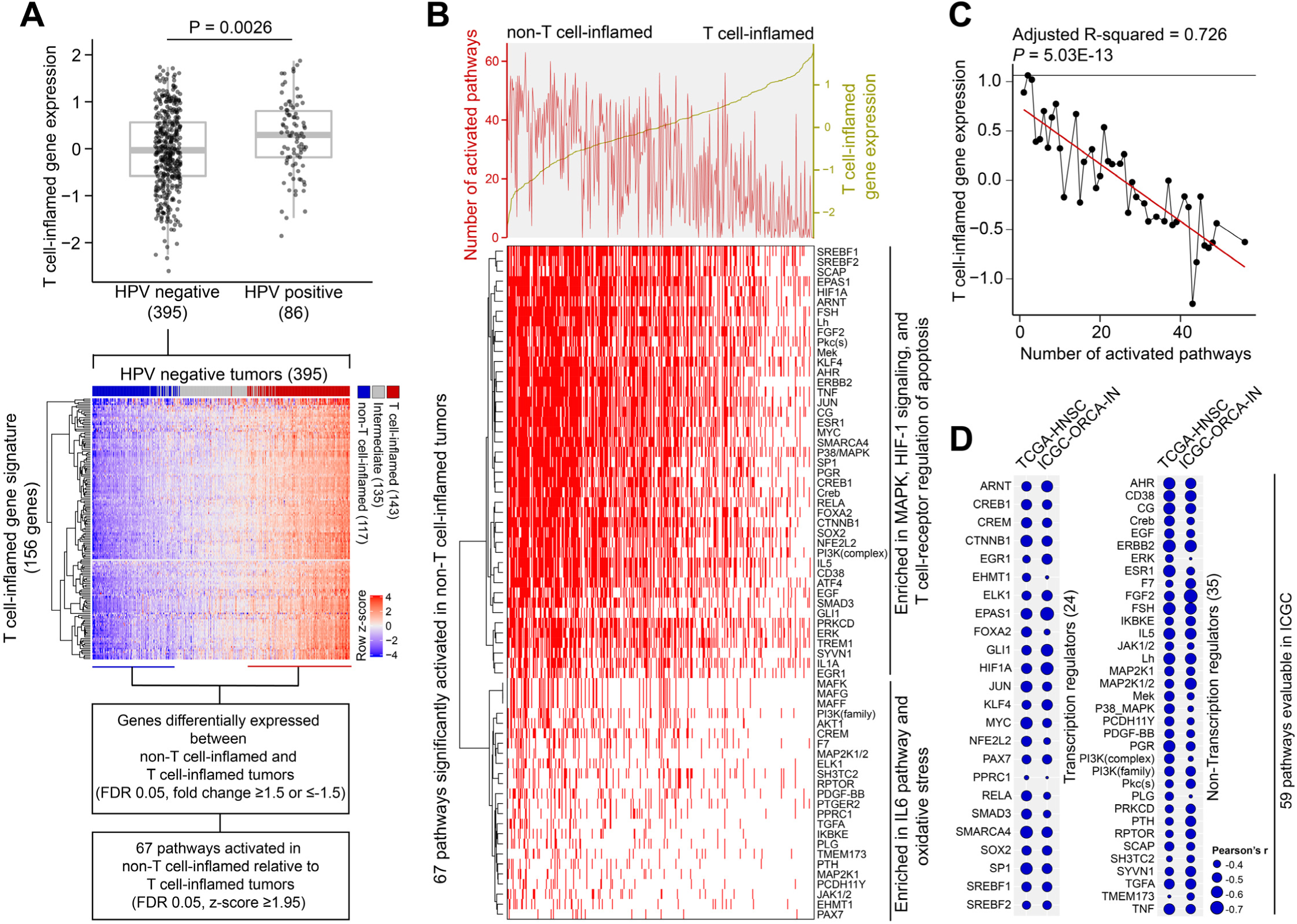
Activation of pathways associated with non-T cell-inflamed tumor microenvironment in HPV-negative head and neck squamous cell carcinoma (HNSCC). (A) Analysis workflow to identify pathway activation in non-T cell-inflamed relative to T cell-inflamed tumors using bulk RNAseq from TCGA. In the boxplot panel, the median is shown by the horizontal center line, with the 25th and 75th percentiles depicted by the boxes; the whiskers extend to the outermost data point within a range of up to 1.5 times the interquartile range (IQR); each data point represents one tumor sample. (B) Landscape of pathway activation in tumor (*n*=395) identified using causal network analysis from Ingenuity Pathway Analysis (IPA). On the heatmap, rows represent the immune-exclusion pathways, and columns represent the tumor specimens. From left to right, samples were sorted by lower to higher T cell-inflamed gene expression. On the dual-axis chart above the heatmap, the primary *y*-axis on the left denotes the number of activated pathways in a sample, and the secondary *y*-axis on the right denotes T cell-inflamed gene expression per sample. (C) Reduced T-cell inflamed gene expression in tumors correlates with a higher number of co-occurring activated pathways. *y*-axis denotes the pathway scores same as in C. *x*-axis denotes T cell-inflamed gene expression averaged across all tumors with the same pathway score. (D) ICGC validation of activated pathways discovered in TCGA. For boxplot in A and line plot in C, unit on the y-axis is log2-transformed and normalized read counts from RNAseq data. Two-sided Welch Two-Sample *t*-test was used in comparing HPV groups (A, upper panel), limma voom with precision weights was used in identifying differentially expressed genes (A, first text box), linear regression was used in C, Pearson’s correlation was used in D. P-values shown are after Benjamini-Hochberg (BH) FDR adjustment for multiple comparisons.

Consistent with our previous pan-cancer analysis(12), the number of co-occurring activated immune-exclusion pathways per tumor was significantly and inversely correlated with T cell-inflamed gene expression in HPV-negative HNSCC (R^2^=0.726, *P*<0.05) (**Fig. 1C**). Of 67 pathways, 59 were independently validated in an oral cancer cohort from The International Cancer Genome Consortium(24) (ICGC-ORCA; *P*<0.05) (**Fig. 1D**), with 24 driven by transcriptional factors such as HIF1A, and the remaining 35 by kinases, hormones, or other non-transcriptional factors (**Fig. 1D**). By cross-referencing to existing drug databases including FDA-approved agents, we identified a catalog of novel molecular targets associated with the non-T cell-inflamed tumor that are ripe for potential drug repurposing (**Table S1**).

### Multimodality analysis identified tumor cell-specific p38 MAPK activation as the master regulator linked to low T cell infiltration in the TME

To investigate the cellular source of these immune-exclusion pathways, we analyzed two independent HPV-negative HNSCC single-cell(sc) RNAseq datasets (cohort #1: Puram et al.(25); cohort #2: Kurten et al.(26)), selecting advanced stage (T3/4) primary tumors. In cohort #1 (2,435 cells from 11 patients), UMAP analysis resolved distinct populations including epithelial cells, fibroblasts, T cells, B cells, and macrophages (**Fig. 2A**). For each pathway, we computed an expression score in individual cells by averaging the expression of downstream target molecules. Among all cell populations, malignant epithelial cells showed the highest pathway scores, suggesting that tumor cells are the major contributor to the immune-exclusion pathway expression in TME, consistent with our hypothesis (**Fig. 2B**).

**Figure 2.**
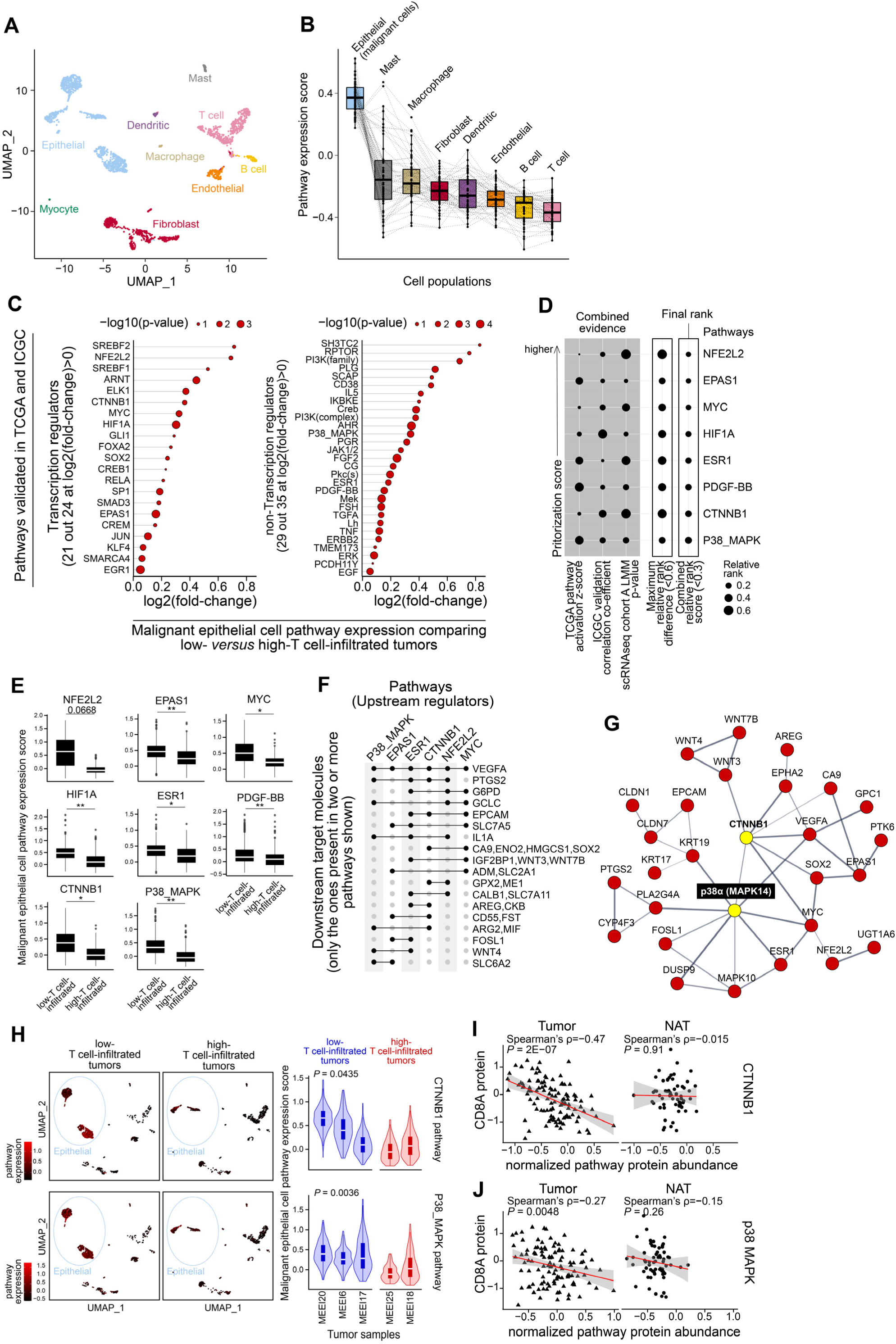
Immune-exclusion oncogenic pathways are activated in malignant cells from HPV-negative head and neck squamous cell carcinoma (HNSCC) of low T cell infiltration. (**A-H**) HNSCC scRNAseq cohort #1 (Puram et al.). (**A**) Distribution of tumor, stroma, and immune cell subsets on UMAP (n=2,435 cells from 11 tumors). (**B**) Expression of 59 pathways from Fig. 1D across cell populations. Pathway scores were computed as the average expression of all genes involved in a pathway. Each data dot is the average score of a pathway in each cell population across all samples. Dotted lines connect the per-cell population scores of the same pathway. (**C**) 50 out of 59 pathways showed higher expression in 774 malignant epithelial cells of low-T cell-filtrated relative to 253 malignant epithelial cells of high-T cell-infiltrated tumors. (**D**) Representative eight pathways that passed prioritization score (combined relative rank) < 0.6 and maximum relative rank difference < 0.3. For each of the 59 pathways from Fig. 1D, its combined relative rank was computed as the geometric mean of three values: the relative rank in TCGA pathway activation (z-score higher to lower), the relative rank in ICGC anti-correlation with T cell-inflamed expression (coefficient highly to lowly negative), and the relative rank in scRNAseq pathway expression comparing malignant epithelial cells from low-T cell-infiltrated *versus* high-T cell-infiltrated tumors (p-values smaller to larger). (**E**) Expression of the eight pathways in the same order as shown in **D**. Denotation: ** FDR-adjusted *P*<0.01, * FDR-adjusted *P*<0.05, otherwise the numbers are shown. (**F**) Six out of eight oncogenic pathways strongly associated with a non-T cell-inflamed TME share downstream target molecules from curated literature in Ingenuity Knowledge Base. (**G**) Protein-protein functional interaction network based on the six pathways and shared downstream targets from **F**. Nodes with at least one connection are shown, from STRING functional protein association networks (confidence score >0.4; active interaction sources as “Experiments”, “Databases”, and “Co-expression”). (**H**) Expression of CTNNB1 pathway and p38 pathway from low-*versus* high-T cell-infiltrated tumors. Five out of 11 tumors with at least 40 malignant epithelial cells per sample were included in analysis. (**I-J**) Proteomics correlation between CD8A protein abundance and (**I**) CTNNB1 or (**J**) p38 MAPK pathway proteomic score in HPV-negative HNSCC specimens from the CPTAC database. Tumors and adjacent normal tissues (NAT) were shown. Pathway score was computed by averaging the normalized protein abundance of target molecules as defined in Ingenuity Knowledge Base. For boxplots in **B**, **E**, and **H**, the median is shown by the horizontal center line, with the 25th and 75th percentiles depicted by the boxes; the whiskers extend to the outermost data point within a range of up to 1.5 times the interquartile range (IQR); unit on the y-axis is scaled log_2_(TPM/10+1) from scRNAseq data. Linear mixed-effects model via maximum likelihood was used in **C**, **E,** and **H**, with tumor group as the fixed effect and tumor id as the random effect. Likelihood ratio test (LRT) was used with the fitted model for computing p-values, followed by BH-FDR adjustment for multiple comparisons. Spearman’s correlation was used in **I-J**.

Next, we sought to understand differences in tumor cell-specific pathway expression for the low-*versus* high-T cell-infiltrated groups. For initial clarity, we focused on the phenotypes based on T cell fraction out of all cells sequenced in the TME per sample (<20%: low infiltration; ≥20%: high infiltration; based on previous work(26)). In cohort #1, 49 out of 59 pathways showed significantly elevated expression in malignant epithelial cells from tumors with low T cell infiltration (FDR-adjusted *P*<0.10) (**Fig. 2C**; **Table S2**). These findings were independently validated cohort #2 (**Fig. S2A-S2Ca**; **Table S3**). For cohort #2 we had tissue access for four samples (26) and the T cell high/low infiltration threshold (20%) was verified by pathologist-evaluated lymphocyte-based inflammation scores from the same tumors (**Table S4**). Significant enrichment of pathway expression was detected exclusively in tumor cells and dominantly in low-T cell-infiltrated tumors across both cohorts, supporting a tumor cell-intrinsic role for those oncogenic pathways in regulating immune exclusion.

To prioritize novel druggable targets for potential combination immunotherapy, we developed a composite scoring system integrating pathway activation data from TCGA, anti-correlation with T cell-inflamed expression in ICGC, and scRNAseq tumor cell pathway scores across both HNSCC cohorts (**Fig. 2D, 2E**; **Fig. S2D, S2E**; **Table S5**). This approach narrowed the 49 candidates to six pathways (**Fig. 2F**), noting that VEGFA, a known mediator of ICI resistance(27), is a downstream target shared by all pathways (**Fig. 2F**). Further, these pathways form a core biomolecular protein-protein interaction network in non-T cell-inflamed tumors centered at p38 (MAPK14) and CTNNB1, with additional nodes including NFE2L2, EPAS1, ESR1, and MYC (**Fig. 2G**). Tumor cell expression of both CTNNB1 and p38 pathways was significantly upregulated in low-relative to high-T cell-infiltrated tumors in both scRNAseq cohorts (FDR-adjusted *P*<0.05; **Fig. 2H**; **Fig. S2F**).

We next validated CTNNB1 and p38-activated tumors at the protein level in high-throughput tumor proteomics data of 101 HNSCC HPV-negative tumors from Clinical Proteomic Tumor Analysis Consortium(28) (CPTAC). We observed an inverse correlation between CD8A protein abundance and CTNNB1 or p38 pathway target molecules (Spearman’s ρ=−0.47 and −0.27, respectively) (**Fig. 2I, 2J**). This inverse correlation was only detected in tumor tissues (*P*<0.01) and not in adjacent normal tissue (NAT) (*P*>0.25), further supporting that activation of CTNNB1 and p38 causally leads to immune exclusion in some tumors. To orthogonally validate these findings at the protein level in human tissues, we performed immunohistochemistry (IHC) on 68 cores from tissue microarrays (TMA) of oral cavity HNSCC. We focused on phosphorylated p38 (phospho-p38), the active form of the kinase(14, 15). Phospho-p38^+^ and CTNNB1^+^ cell densities were positively correlated (Spearman’s ρ=0.30, *P*=0.014), and in CTNNB1-high tumors, phospho-p38-high cores had significantly fewer CD8^+^ TILs than phospho-p38-low cores (*P*=0.038) (**Fig. S3A-B; Table S6**).

Between the two candidates, we elected to focus on p38 for further investigation. Consistent with our prior work(4, 5, 12, 21), CTNNB1 was strongly associated with non-T cell-inflamed tumors, however the development of therapeutics to target CTNNB1 pharmacologically has been challenging due to toxic and poorly effective drugs(29). In contrast, a tumor cell-intrinsic role for p38 in immune exclusion has not been previously described, and multiple small-molecule p38 inhibitors have advanced to clinical testing, making it a tractable candidate for therapeutic intervention(30, 31). Taken together, through a rigorous multi-step prioritization strategy integrating bulk RNAseq, scRNAseq, and proteomics data across independent cohorts (**Fig. S4**), we identified p38 MAPK (MAPK14) as a central hub in the immune-exclusion pathway network. Noting the lack of tolerable, high-potency inhibitors of CTNNB1(18), we focused on p38 given the availability of multiple clinical-stage inhibitors. Tumor cells expressed the highest levels of p38 pathway activation compared to non-epithelial cells including fibroblasts (**Fig. S5**), further supporting a tumor-intrinsic role for p38 as a high-priority candidate for immune-oncology translation.

### Multi-cancer validation of tumor cell p38 activation in association with low T cell infiltration across tumor types

Our prior work demonstrated CTNNB1 activation as driving the absence of T cell infiltrates in multiple tumor types(4, 5), and we were interested in whether p38 may be a similar broad mechanism of immune exclusion. To investigate this, we generated a 12-gene p38 pathway activation signature by an integrative analysis of multi-omics data from TCGA, ICGC, scRNAseq, and CPTAC (*ARG2, CD55, CYP4F3, FST, GCLC, IL1A, MIF, PLA2G4A, PTGS2, S100A12, SLC6A2, VEGFA*), and shifted our analysis to LUSC, a tumor type of very similar histology and molecular biology as HNSCC(20, 32, 33). We repeated the same analysis in two LUSC scRNAseq cohorts (cohort #3: Mu et al.(34); cohort #4: Qian et al.(35)). We validated our previous findings in both cohorts, including enrichment of p38 MAPK pathway expression in malignant epithelial *versus* other cells (**Fig. 3A-B**) or fibroblasts specifically (**Fig. 3C-D**), and significantly higher p38 MAPK pathway expression in tumor cells from low-*versus* high-T cell-infiltrated tumors (*P*=0.00019 in **Fig. 3E**; 2.58E-06 in **Fig. 3F**). We then expanded these scRNAseq analyses to cohorts of other tumor types known to be responsive to anti-PD1, including lung adenocarcinoma (LUAD; cohort #5(36)), renal clear cell carcinoma (RCC, cohort #6(37)), and skin cutaneous melanoma (SKCM, cohort #7(38)). Comparing low-*versus* high-T cell-infiltrated tumors, we identified a significantly higher tumor cell-expressing p38 activation score in low-T cell-infiltrated tumor specimens from LUAD and RCC (*P*<0.05; **Fig. 3G-H**), but not in the SKCM scRNAseq cohort (**Fig. 3I**), suggesting that certain tumor cell-intrinsic signaling pathways may impact immunotherapy outcomes in different cancers.

**Figure 3.**
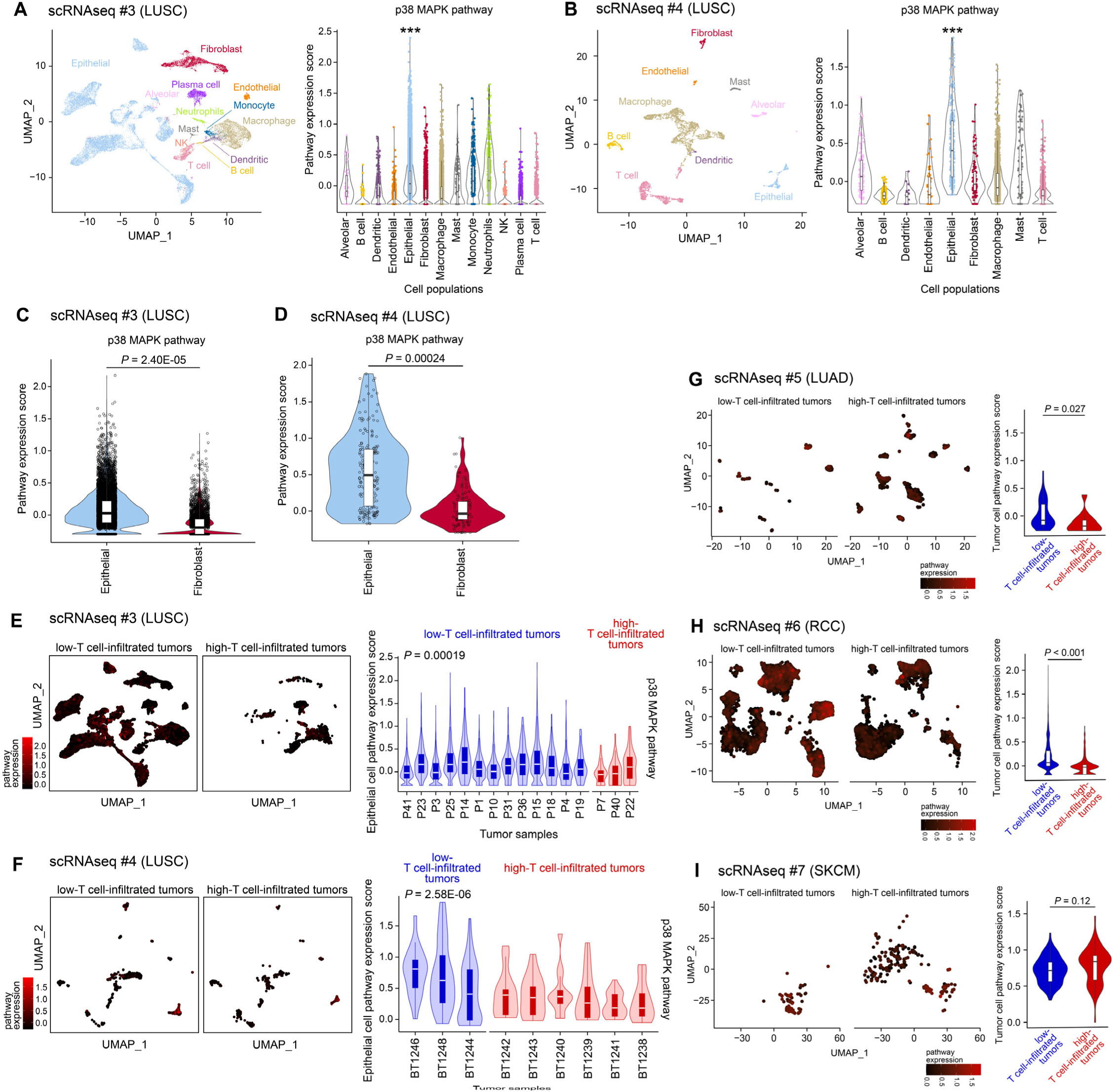
**Validation of tumor cell p38 pathway association with low T cell infiltration across solid tumors**. Two scRNAseq cohorts (#3, #4) of lung squamous cell carcinoma (LUSC) and three cohorts (#5, #6, #7) of lung adenocarcinoma (LUAD), renal clear cell carcinoma (RCC), and cutaneous skin melanoma (SKCM) are shown. (**A-B**) Distribution of tumor, stroma, and immune cell subsets on UMAP, and expression of p38 MAPK pathway across cell populations in LUSC. scRNAseq #3 (LUSC) in **A**: n=28,322 cells from 16 tumors. scRNAseq #4 (LUSC) in **B**: n=3,444 cells from ten tumors. Pathway scores were computed as the average expression of all genes involved in a pathway. Denotation: *** *P*<0.0001. (**C-D**) p38 pathway expression scores in malignant epithelial cells and fibroblasts in LUSC. (**E-F**) Expression of p38 pathway in low-*versus* high-T cell-infiltrated tumors in LUSC. scRNAseq #3 (LUSC) in **E**: n=17,169 malignant epithelial cells total from 13 low-T cell-infiltrated tumors and 164 malignant epithelial cells total from three high-T cell-infiltrated tumors are shown, respectively. scRNAseq #4 (LUSC) in **F**: n=155 malignant epithelial cells total from three low-T cell-infiltrated tumors and 92 malignant epithelial cells total from six high-T cell-infiltrated tumors are shown, respectively; one tumor with less than 10 epithelial cells was excluded from this comparison. (**G-I**) Single-cell analysis of p38/MAPK activation across tumor types of LUAD, RCC, and SKCM. P-value was computed by linear mixed-effects models (LMM) with tumor group as the fixed effect and tumor id as the random effect, followed by likelihood ratio test (LRT) in **A-F**. A nested LMM design (cell type nested within tumor id) was used in **A-D**. Two-sided Welch Two Sample *t*-test was used in in **G-I**. For boxplots in **A-I**, the median is shown by the horizontal center line, with the 25th and 75th percentiles depicted by the boxes; the whiskers extend to the outermost data point within a range of up to 1.5 times the interquartile range (IQR).

### p38 MAPK driving the non-T cell-inflamed TME validated by multispectral imaging and spatial transcriptomics in situ

Having established the association between tumor cell p38 activation and reduced T cell infiltration computationally across multiple tumor types, we next sought to validate these findings in human tumor tissues using orthogonal spatial approaches. Using multispectral immunofluorescence (mIF) imaging with PhenoImager^TM^ HT on HNSCC and LUSC tumor TMAs, we showed that PanCK^+^ tumor cells had significantly higher abundance of phospho-p38 relative to CD3^+^ T cells, CD68^+^ macrophages, and other cells present in the TME, supporting our findings above that tumor cells are the main cellular source for p38 pathway activation in the TME (**Fig. S6A-D**).

We next investigated the spatial relationship of phosphorylated p38^+^ tumor cells and CD8^+^ TILs on 154 HN oral cavity cores (105 primary tumors and 49 normal mucosa; Panel markers: PanCK, CD3, CD8, PD1, phospho-p38 plus DAPI). Tumor-infiltrating CD3^+^CD8^+^ T cells, particularly the CD3^+^CD8^+^PD1^+^ subset, were depleted in tumors with high phospho-p38 relative to those with low phospho-p38 (**Fig. 4A**; **Table S7**). Phospho-p38^+^PanCK^+^ cell density was significantly and inversely correlated with CD3^+^CD8^+^ T cells (Spearman’s ρ=−0.43), and CD3^+^CD8^+^PD1^+^ T cells (ρ=−0.37), a pattern only detected in tumors but not in normal mucosa (*P*<0.05; **Fig. 4B**). In cases where multiple cores were derived from the same patient, we repeated the analysis after averaging cell densities to the patient level, and the inverse correlation remained significant for tumor specimens (ρ=−0.57 for both; *P*<0.05). To gain insights into the spatially resolved gene expression changes associated with phospho-p38, we performed digital spatial profiling (DSP GeoMX) on 32 regions of interest (ROIs) in tumor cores from the HNSCC tissue cohort. Tumor cells within each ROI were segmented by PanCK^+^ morphology marker staining (**Fig. 4C**). Comparing phospho-p38 high *versus* low tumor cells (**Fig. 4D**), 141 downregulated and 324 upregulated differentially expressed genes (DEGs) were detected at fold change ≤-1.5 or ≥ 1.5, respectively (FDR-adjusted *P*<0.10; **Fig. 4E**; **Table S8**). Pathway analysis revealed that the downregulated genes in phospho-p38 high tumor cells were significantly enriched in IFN signaling regulation, and upregulated genes were enriched in MYC signaling and fatty acid metabolism, among others (FDR-adjusted *P*<0.10; top pathways shown in **Fig. 4F**; full result provided in **Table S9**).

**Figure 4.**
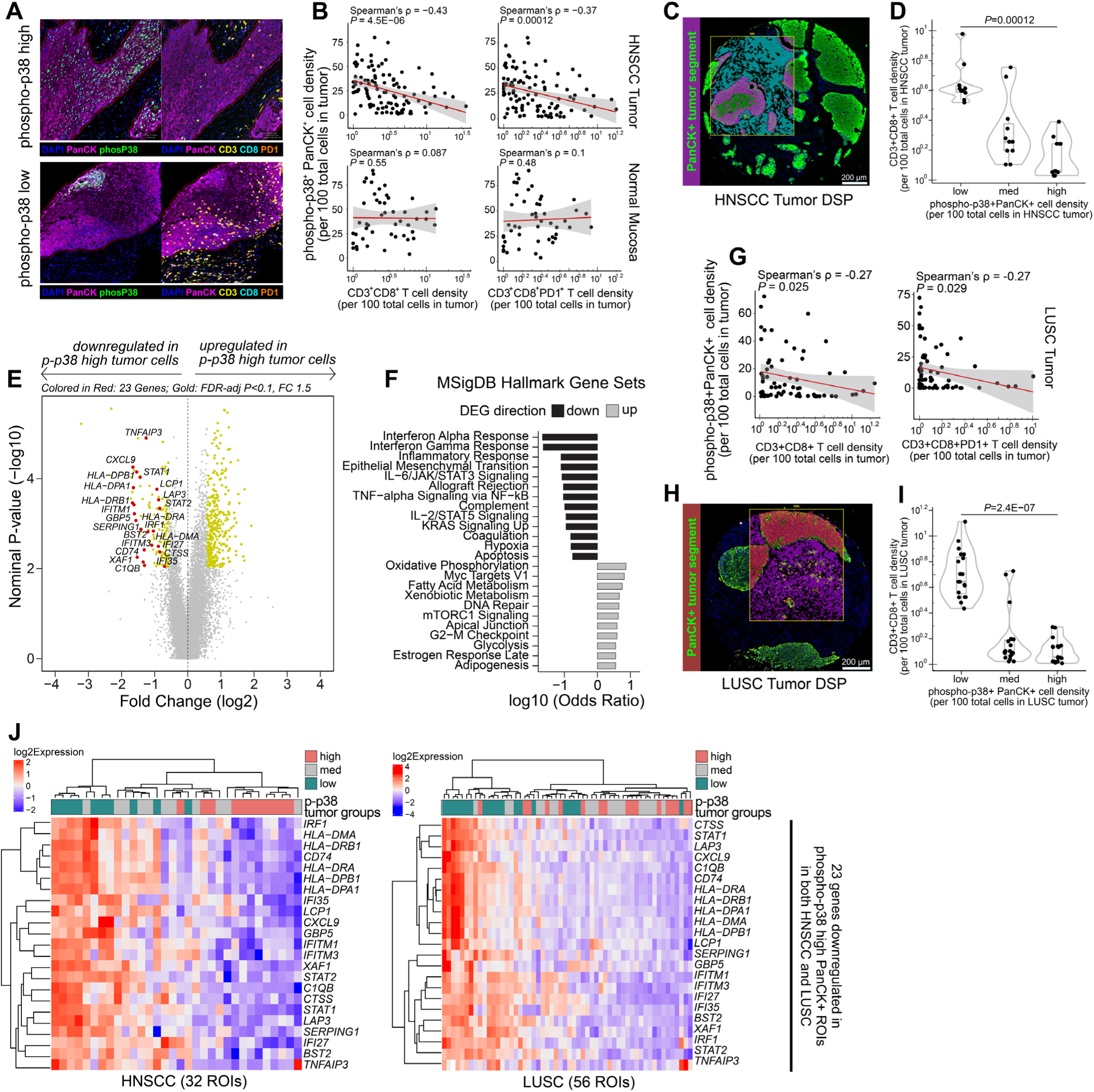
p38 MAPK pathway in non-T cell-inflamed tumors by multispectral immunofluorescence (mIF) imaging and spatial transcriptomics. (**A**) mIF imaging of phospho-p38^+^ high and low tumors showing the CD3^+^CD8^+^ and CD3^+^CD8^+^PD1^+^ T cell infiltrates in PanCK^+^ tumor compartment in HNSCC. (**B**) Correlation between phospho-p38^+^PanCK^+^ cell density and CD3^+^CD8^+^ T cell, CD3^+^CD8^+^PD1^+^ T cell in HNSCC tumor (n=105 cores) or normal mucosa (n=49 cores). (**C**) PanCK^+^ segmentation mask (denoting tumor cells) from digital spatial profiling (DSP) of HNSCC tumor cores. Box represents selected regions of interest (ROI; max size 660×785 µm). Inside each ROI, PanCK^+^ segmentation is labeled as dark purple, with rest as cyan. (**D**) High/med/low phospho-p38 HNSCC tumor cores selected for DSP experiment (n=32). Tertiles were used to segregate tumors into three groups based on phospho-p38^+^PanCK^+^ cell density from multispectral imaging analysis: low (n=10), med (n=11), and high (n=11). (**E**) DEGs comparing tumor cell segment of high/low phospho-p38 HNSCC DSP from **D**. p-p38 = phospho-p38. (**F**) Pathway enrichment of down and upregulated DEGs from **E**, using Enrichr(61, 62) with MSigDB hallmark 2020 gene sets. (**G**) Correlation between phospho-p38^+^PanCK^+^ cell density and CD3^+^CD8^+^ T cell, CD3^+^CD8^+^PD1^+^ T cell in LUSC tumor (n=68 cores). (**H**) PanCK^+^ segmentation mask (denoting tumor cells) from DSP of LUSC tumor cores. Inside each ROI, PanCK^+^ segmentation is labeled as dark red, with rest as purple. (**I**) High/med/low phospho-p38 LUSC tumor cores selected for DSP experiment (n=56). Tertiles were used to segregate tumors into three groups based on phospho-p38^+^PanCK^+^ cell density from multispectral imaging analysis: low (n=20), med (n=19), and high (n=17). (**J**) Heatmaps of 23 downregulated DEGs from HNSCC tumor DSP (labeled in **E**) validated in LUSC (FDR-adjusted *P*<0.10). For boxplots in **D** and **I**, the median is shown by the horizontal center line, with the 25th and 75th percentiles depicted by the boxes; the whiskers extend to the outermost data point within a range of up to 1.5 times the interquartile range (IQR). Spearman’s correlation was used in **B** and **G**, two-sided Wilcoxon rank sum test in **D** and **I,** two-sided Fisher’s exact test in **F**. Linear mixed-effects model was used in **E** and **J**, with tumor group as the fixed effect and slide id as the random effect. P-values were adjusted by BH-FDR procedure.

Given our findings above that suggested p38 may be associated with immune exclusion across epithelial cancer types, we repeated the same tissue validation experiment in LUSC consisting of 68 primary tumors. Similar to HNSCC, phospho-p38^+^PanCK^+^ cell density was significantly and inversely correlated with CD3^+^CD8^+^ T cells and CD3^+^CD8^+^PD1^+^ T cells (Spearman’s ρ=−0.27 for both, *P*<0.05; **Fig. 4G**; **Table S10**). To validate the down-regulated DEGs identified from HNSCC DSP in LUSC, we performed DSP on 56 ROIs in LUSC tumor cores followed by PanCK^+^ segmentation for tumor cells (**Fig. 4H**). Comparing phospho-p38 high *versus* low tumor cells (**Fig. 4I**), 23 genes were consistently downregulated in phospho-p38 high PanCK^+^ ROIs in both HNSCC and LUSC tumor types (**Fig. 4J**; **Table S11**). These notably included T cell chemokines (i.e. - CXCL9), and MHCII genes, representative of antigen-representation machinery. Taken together, these in situ tissue validation experiments support our in silico discovery that tumor cell p38 MAPK activation is associated with immune exclusion.

### p38 inhibition in human cancer cell lines increases T cell recruitment in vitro

To functionally investigate the impact of tumor cell p38 inhibition on T cells, we pursued a series of in vitro experiments that may provide further mechanistic insights into p38 inhibition as an immune-modulatory mechanism, using two human HPV-negative HNSCC (PCI13, PCI4B) and two LUSC (H520, H1703) cell lines, using both genetic and pharmacologic approaches.

We first established shRNA knockdown (KD) of *MAPK14* (encoding p38α) in all four cancer cell lines, with protein confirmation by Western Blot (WB) (**Fig. 5A**), and transcript confirmation by RT-PCR (**Fig. 5B**). We collected conditioned media from p38-KD and wildtype (WT) cultures (empty vector [EV] controls), and demonstrated that T cell migration was increased upon co-culture with conditioned media from p38-KD *versus* WT cancer cell lines (**Fig. 5C**, by Transwell assays). To detect expression changes induced by p38-KD in cancer cells, we performed RNAseq and identified DEGs in each cell line. Among significant DEGs (FDR-adjusted *P*<0.05, fold change ≥1.1), 11 genes showed consistent expression increases across four cell lines in the p38-KD group (**Fig. 5D**). This includes *CXCL16*, a prominent cytokine that has been previously reported to attract T cells via CXCL16-CXCR6 axis and associate with better survival in patients with cancer(39–41) (**Fig. 5E**). We orthogonally validated elevation in *CXCL16* transcripts in cancer cell lysates using RT-PCR (**Fig. 5F**), and its protein in cancer cell-conditioned media using ELISA (**Fig. 5G**).

**Figure 5.**
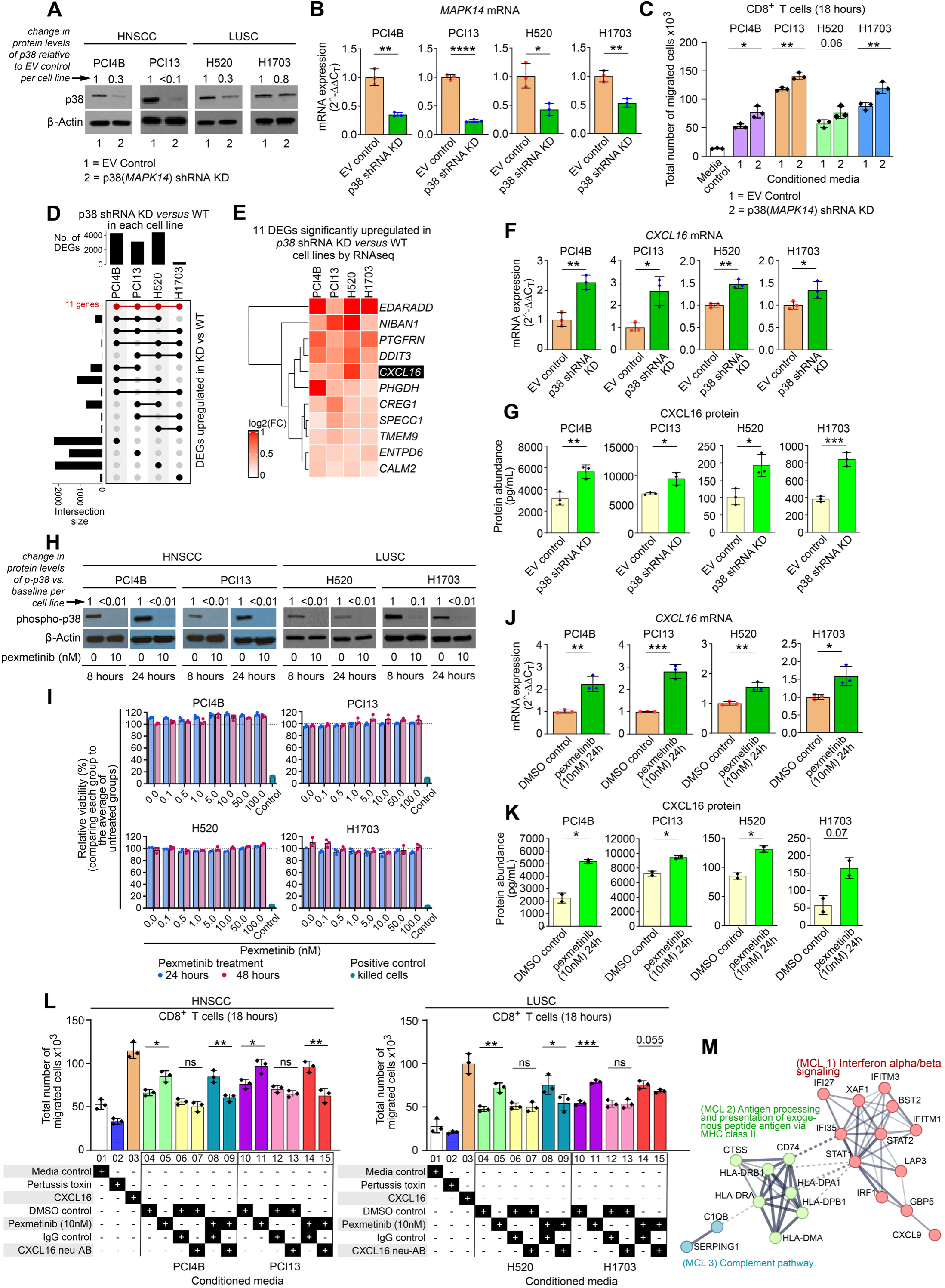
p38 inhibition in human cancer cell lines enhances CD8⁺ T cell recruitment in head and neck squamous carcinoma (HNSCC) and lung squamous cell carcinoma (LUSC). (A-G) p38α (encoded by *MAPK14*) shRNA knockdown (KD). (**H-L**) pharmacological p38 inhibition by pexmetinib. (**A**) Western blot analysis of p38α expression in p38 KD and EV (empty vector) control cells from human HNSCC cell lines PCI13 and PCI4B, and LUSC cell lines H520 and H1703. ag. (**B**) Quantification of p38 KD efficiency by real-time(RT) PCR, showing 50% or more reduction in p38α expression in p38 KD cells compared to EV controls. (**C**) Migration of CD8^+^ T cells (from healthy donor) toward conditioned media from EV and p38 KD cells in HNSCC and LUSC cell lines. (**D**) 11 genes were consistently upregulated in p38 KD *versus* WT (EV controls) RNAseq across four cell lines shown as UpSet plot. (**E**) Gene expression heatmap of the 11 genes from **D**. Black box highlights CXCL16, a chemokine known to attract T cells. (**F**) RT-PCR analysis validates the upregulation of CXCL16 mRNA in p38 KD cells compared to EV controls. (**G**) ELISA assay validates increased CXCL16 protein release into the conditioned media from p38 KD cells compared to EV controls. (**H**) Western blot confirming inhibition of p38 phosphorylation in HNSCC and LUSC cell lines treated with the p38 inhibitor pexmetinib at 10 nM (8 and 24 hours). (**I**) Cell viability is not reduced upon increasing pexmetinib dose by CellTiter-Glo^®^ luminescent assay, which quantifies intracellular ATP as a proxy for metabolically active (viable) cells (24 and 48 hours). *y*-axis indicates relative viability of each group normalized against the average of all untreated groups (0 nM pexmetinib) for robustness. (**J**) RT-PCR analysis validates upregulation of CXCL16 mRNA levels in pexmetinib-treated cancer cells, similar to p38 KD cells. (**K**) ELISA assay validates increased CXCL16 protein release into conditioned media from pexmetinib-treated cancer cells. (**L**) CD8⁺ T cell migration assay demonstrates increased migration toward conditioned media from pexmetinib-treated HNSCC (PCI13 and PCI4B; **left**) and LUSC (H520 and H1703; **right**) cells, confirming enhanced recruitment. The addition of a CXCL16-blocking antibody to the conditioned media significantly reduced CD8⁺ T cell migration, indicating that CXCL16 plays a key role in mediating T cell recruitment in response to tumor cell p38 inhibition. (**M**) Protein-protein functional interaction network based on the 21 genes from **J** (two unconnected genes not shown), using STRING functional protein association networks with nodes colored by Markov Cluster Algorithm(70) (MCL) clusters. In **A-C, F-G, H-L**, each experiment was performed at least twice; representative data from one experiment are presented as mean ± SD. Two-sided unpaired Student *t*-test was used in **B-C**, **F-G**, **J-L**. P-values were computed comparing groups of interest within each cell line, without multiple comparisons across cell lines. Denotation: **** *P*<0.0001, *** *P*<0.001, ** *P*<0.01, * *P* < 0.05, exact p-values shown if *P*<0.10. ns = not significant. FDR was controlled at 0.10.

In addition to p38-KD, we examined the trans-impact of pexmetinib, a selective small molecule p38 inhibitor that has been evaluated clinically (30, 31) (also known as ARRY-614; see **Methods**), on tumor cell-mediated T cell migration in vitro. We first confirmed that treatment of cancer cells with pexmetinib inhibited p38 phosphorylation (**Fig. 5H**, by WB). Tumor cell viability was not affected by increasing doses of pexmetinib, suggesting pexmetinib does not kill tumor cells directly (**Fig. 5I**). This was accompanied by increases in *CXCL16* transcript levels in cancer cells (**Fig. 5J**, by RT-PCR) and CXCL16 protein release into the media (**Fig. 5K**, by ELISA). This finding was consistent with the p38-KD cancer cell results described above. We collected conditioned media from pexmetinib-treated and DMSO control cancer cells, removed pexmetinib by a 3 kDa Amicon^®^ Ultra Centrifugal Filter (Millipore Sigma; retaining proteins while discarding small molecules), and then co-cultured the media with T cells. We demonstrated that T cell migration increased upon co-culture with conditioned media from pexmetinib-treated tumor cells *versus* control, similar to our observations in p38-KD experiments above. Of note, neutralizing CXCL16 by a human/primate CXCL16 antibody abrogated the increase in T cell migration, suggesting that the effect is CXCL16-mediated (**Fig. 5L**). To ensure accuracy, we took extra caution in removing the drug substance after tumor cell treatment and before co-culturing the conditioned media with T cells, and confirmed that neutralization of CXCL16 protein was successful by ELISA. Taken together, these findings provide mechanistic support that p38 inhibition in tumor cells increases cytokine release, with CXCL16 as an example in these cancer cell lines and enhances T-cell migration.

Observing this confirmation of a connection between p38 and T cell chemokine gradients, we returned to the tumor-based DSP experiments to further investigate DEGs in human tissues. By protein-protein interaction network analysis from STRING(42), we found that the 23 genes consistently down-regulated in high phospho-p38 tumor cells in HNSCC or LUSC form two main clusters representing IFNα/β signaling (cluster 1) and MHC II antigen processing and presentation (cluster 2) (**Fig. 5M**). Noting gene expression variance between cancer cell lines and human tumor tissues, these findings provide consistent evidence supporting a conceptual model wherein p38 inhibition releases tumor cell suppression of chemokine gradients, which in turn facilitates T cell infiltration in the TME.

## DISCUSSION

Although p38 MAPK has been implicated in cellular stress responses across multiple cell types, we report here a previously unrecognized, tumor cell-intrinsic role for p38 activation in promoting immune exclusion. While prior studies have described p38 function in stromal and immune cells, our comprehensive computational and experimental analyses - spanning *in silico* modeling, proteomics, spatial transcriptomics, and *in situ* validation - identify tumor cells as the predominant source of active p38 signaling in the TME. Functional assays demonstrate that p38 inhibition, using both shRNA and small-molecule inhibitor, restores chemokine signaling (e.g., CXCL16) and enhances T cell recruitment.

While existing biomarkers for immune checkpoint inhibitor (ICI) responsiveness, such as PD-L1 expression and TMB, focus on T cell infiltration or neoantigenicity, a majority of tumors do not meet these criteria and remain refractory to ICIs. The non-T cell-inflamed phenotype, characterized by minimal IFN signaling, impaired antigen-presenting cell (APC) activity, and low cytolytic marker expression, provides a conceptual framework to understand resistance to immunotherapy. Oncogenic pathways such as WNT and MYC programs have been linked to immune exclusion, yet efforts to therapeutically target these pathways have often been hindered by toxicity, limited efficacy, or unintended immune suppression(43). Through a discovery-validation framework, we have identified functional crosstalk between p38 MAPK and these oncogenic programs. Importantly, p38 inhibition offers a clinically tractable, tolerable approach for modulating the non-T cell-inflamed TME. Beyond p38, we have generated a catalog of candidate targets and pharmacological agents for drug repurposing, based on the hypothesis surrounding their capacity to reverse immune suppression and release chemokine gradients to facilitate anti-tumor responses.

An important nuance in our findings is that the specific chemokine mediators identified differ between experimental systems. We found that p38 inhibition upregulated CXCL16 in human cancer cell lines, whereas in human tumor tissues, malignant epithelial cells from phospho-p38-low tumors expressed higher levels of CXCL9 and MHC class II antigen-presentation genes compared to those from phospho-p38-high tumors. This likely reflects the broader immunomodulatory program regulated by tumor cell p38 in the complex tissue microenvironment, where CXCL9 expression is IFNγ-dependent(44, 45) and thus not captured in cell line cultures. Despite these context-dependent differences, the findings converge on a consistent model, where p38 inhibition in tumor cells enhances CD8^+^ T cell recruitment by upregulating chemokine expression. Indeed, previous studies reported that cancer cells are major sources of CXCL16 and IFNγ-inducible CXCL9, each attracting TILs and are associated with better prognosis in lung, colorectal, breast, and ovarian cancers(39, 45–47).

While we emphasize here the tumor cell-intrinsic role of p38 in immune exclusion, we would also note that p38 plays complex and context-dependent roles in other cell types within the TME. In breast cancer stroma, ablation of p38 led to M1-like macrophage differentiation and CD4^+^ T cell infiltration(19). In macrophages, p38 MAPK activation regulates polarization toward the immunosuppressive M2-like state(48), consistent with the increased density of M2-like macrophages generally associated with non-T cell-inflamed tumors(49) and moreover underlies interest in p38 inhibitors outside of oncology such as chronic inflammatory and autoimmune disorders. In DCs, p38 has been described to promote maturation and costimulation(50), however paradoxically also drives a tolerogenic DC state associated with tumor formation(51). Additionally, p38 inhibition during early DC differentiation enables an OX40 dependent DC program to activate effector T cells in tumor-specific fashion and inhibit regulatory T cell activity(17). Regarding T cells, p38 may play multiple roles in stress responses but notably, in a CRISPR-Cas9-based genetic screen of primary T cells, p38 knockout maximized cell fitness across domains of cell expansion, differentiation as well as oxidative and genomic stress(52). These results further emphasize our robust human biospecimen based discovery and validation approach, taken across HNSCC, NSCLC, RCC, and other tumor types, highlighting the broad relevance of this translational investigation surrounding p38 as a driver of the non-T cell-inflamed TME and as a therapeutic target to overcome immunotherapy resistance across various solid tumors.

We acknowledge that further studies are necessary to fully elucidate the immune-relevant mechanisms underpinning these observations surrounding p38. This study did not characterize signal transduction by which p38 activation in tumor cells leads to suppression of chemokine expression and consequent immune exclusion. While the association between p38 activation and immune exclusion was consistent across epithelial cancer types, it was not observed in a single melanoma scRNAseq cohort, which may reflect tumor lineage-specific differences or limited sample size, and warrants further investigation. While we identified CXCL16 in cell line cultures as a downstream mediator of p38 inhibitor-induced T cell trafficking, future studies incorporating immune-competent ex vivo models and in vivo models will be important to elucidate this therapeutic modality in full.

In summary, using a multi-modality approach spanning bulk and single-cell transcriptomics, proteomics, spatial profiling, and functional validation across multiple epithelial cancer types, we have identified tumor-intrinsic p38 signaling as a driver of immune exclusion that is amenable to therapeutic intervention. These findings provide a rationale for prospective evaluation of p38 inhibition in combination with ICI. More broadly, this work demonstrates a discovery and validation framework and nominates a catalog of candidate targets for novel therapeutic combinations to overcome immunotherapy resistance.

## METHODS AND MATERIALS

### Human specimens

Formalin-fixed, paraffin-embedded (FFPE) tumor tissue samples in the head and neck tissue biobank and lung cancer tissue biobank were obtained from patients consented under The University of Pittsburgh institutional review board (IRB)-approved protocol (HCC#99-069 for head and neck, and STUDY20010250 for lung). All samples have written-informed patient consent.

### Study cohort and datasets

We collected HNSCC cohorts of bulk RNAseq data from TCGA(20) (n=481), ICGC(24) (n=40), and proteomic data from CPTAC(28) (n=101), as well as pan-cancer single-cell sequencing data for the integrative analysis in this study (scRNAseq cohorts #1 to #7). The datasets are from publicly available omics repositories including NCI’s Genomic Data Commons (GDC) and NCBI SRA/GEO with accession IDs listed in key resource table.

### Target prioritization workflow

A hypothesis-driven prioritization strategy was built, providing a strong scientific rationale behind narrowing down p38 MAPK as the final candidate for further validation and clinical translation (**Fig. S4**). *De novo* analysis discovered 67 upstream regulators significantly associated with a non-T cell-inflamed TME using the TCGA-HNSC RNAseq cohort (step 1). The activation of upstream regulators was predicted by Ingenuity Pathway Analysis (IPA)’s causal networks, based on downstream target expression as well as curated Ingenuity Knowledge Base(23). Upstream regulators were referred to as “pathways” throughout the manuscript to distinguish them from single genes or molecules. This initial discovery was followed by multiple validations in independent cohorts, including the ICGC-ORCA RNAseq cohort (step 2), two scRNAseq cohorts (step 3), and integration of all data elements (step 4). These processes narrowed the candidates to six pathways (step 5). Using STRING protein-protein interaction networks(53), CTNNB1 and p38α (MAPK14) were identified as central “hub” nodes (step 6). p38 was chosen for subsequent experimental validation as multiple inhibitors have been previously developed (step 7).

### TCGA RNAseq, ICGC RNAseq, and CPTAC proteomics analysis

In the TCGA cohort, samples were split into HPV-positive and HPV-negative subsets based on HPV status(20). A predefined and validated 160-gene T cell-inflamed gene signature was utilized to categorize tumors into T cell-inflamed, intermediate, or non-T cell-inflamed using a two-tier scoring system following previous protocols established by our group(21). In the ICGC cohort, the T cell-inflamed gene expression was computed by averaging the normalized and log-transformed expression of the 160 genes in the signature and then correlated with each of the oncogenic pathway scores in a continuous manner. For CPTAC, pathway scores of CTNNB1 and p38 MAPK each were computed by averaging the normalized protein abundance of target molecules as defined in Ingenuity Knowledge Base (CTNNB1 pathway: BMP7, CA9, CDH16, ENO2, EPCAM, GAST, GLI2, GPX2, HMGCS1, IGF2BP1, KLF5, MAOA, ME1, PTGS2, SERPINB2, SLC6A2, SOX2, VEGFA, WNT3, WNT4, WNT7B, ZIC5; p38 MAPK pathway: ARG2, CD55, CYP4F3, FST, GCLC, IL1A, MIF, PLA2G4A, PTGS2, S100A12, SLC6A2, VEGFA) and then correlated with CD8A protein abundance.

### Drug-gene interaction

Genes encoding upstream regulators of the 59 oncogenic pathways (e.g., *CTNNB1*, *MAPK14*, etc.) associated with immune exclusion were queried in The Drug Gene Interaction Database(54) (DGIdb, http://www.dgidb.org) (v4.2.0, accessed June 21, 2023) for gene-drug interactions (inhibitor, activator, etc.) with default settings. The DGIdb integrates resources from 40 databases and consists of existing drugs including those that are U.S. Food and Drug Administration (FDA)-approved, antineoplastic, and/or immunotherapies.

### Single-cell RNAseq analysis

For the scRNAseq cancer cohorts, publicly available expression matrices, cell annotations, and malignant cell labels were downloaded and subject to downstream analysis. Within each cohort, the published data were already filtered and normalized from the study, hence no further quality processing steps were needed. For HNSCC cohorts, baseline tumors of HPV-negative and advanced stage (T3/4) were used for analysis. HNSCC cohort #1(25) scRNAseq data were generated using SmartSeq2 and quantified in transcript-per-million (TPM) with log_2_(TPM/10+1) unit, and HNSCC cohort #2(26) scRNAseq data were generated using 10X Genomics 3’ V2 chemistry and quantified in UMI count matrix with log(UMI) unit. Within each dataset, after selecting the first 10 principal components using Principal Component Analysis (PCA), a neighborhood graph was computed based on the PCA representation of the data and visualized by Uniform Manifold Approximation and Projection (UMAP) dimensionality reduction technique with Leiden clustering. For the statistical comparison of tumor cell-expressing pathway scores in low-*versus* high-T cell-infiltrated tumors, data were further filtered to keep tumor samples of ≥40 malignant epithelial cells and ≥100 cells total within each dataset. We repeated the same analysis in two independent LUSC scRNAseq cohorts (Mu et al.(34) & Prazanowska et al.(55), cohort #3; Qian et al.(35) & Gavish et al.(56), cohort #4). For #3, Mu et al.(34) published the cohort initially, and Prazanowska et al.(55) provided more detailed metadata information on the same cohort. For #4, Qian et al.(35) published the cohort initially, and Gavish et al.(56) re-processed the dataset and made it publicly available, which was used in our analysis. Similar analyses were performed for LUAD, RCC, and SKCM scRNAseq cohorts (cohorts #5(36), #6(37), and #7(38)).

### Multi-cancer scRNAseq analysis of p38/MAPK signaling

For each tumor, a p38-activation score was computed as the mean expression of the 12 genes involved in the signature identified in HPV-negative HNSCC, after scaling and centering across all tumor samples. These expression scores were then used to correlate with the T cell-inflamed gene expression across all tumors by Spearman’s correlation. Pathway activation scores were calculated for each tumor sample, requiring at least 50% of the cancer-specific target molecules to be upregulated (relative to its median expression across all samples from an individual cohort) and compared in the non-T cell-inflamed tumor group relative to T cell-inflamed.

### Histopathologic analysis

For histological analysis, 4µm pieces of tumor tissue were fixed in either 10% formalin for H&E staining or Carnoy’s fixative for Periodic-acid Schiff (PAS) staining. Sectioning and staining were performed at University of Pittsburgh Medical Center (UPMC) Translational Oncologic Pathology Services (TOPS). All sections were reviewed by a pathologist in a blinded fashion.

### H&E and CD8, CTNNB1, phospho-p38 immunohistochemistry staining

The hematoxylin and eosin (H&E) and CD8, CTNNB1, phospho-p38 IHC staining on HNSCC FFPE specimens were performed at UPMC TOPS. Slides were cut at 4µm then baked for one hour at 60 degrees Celsius. The slides were cooled to room temperature then deparaffinized and hydrated in diH20. Slides were stained using Hematoxylin 560 MX (3801576), Eosin Phloxine 515 (3801606), Define MX-aq (3803598), and Blue Buffer 8 (3802918) (all catalog numbers from Leica Biosystems). Adjacent sections were used for CD8, CTNNB1, and phosphor-p38 single-plex IHC each. Staining for CD8 (Leica Biosystems, PA0183): the predilute CD8 was stained on the Bond III instrument (Leica Biosystems) for 8 min. Prior to antibody placement, the antigen retrieval used was the ER2 (pH 8.9-9.1, Leica Biosystems, AR9640) for 20 minutes at 95 deg. followed by the Bond Polymer Refine Detection Kit-DAB (Leica Biosystems, DS9800). Staining for phospho-p38 (Cell Signaling, 4511S, 1:50): slides were subjected to deparaffinization and antigen retrieval with CC2 (Roche, 950-223) for 20 mins at 95°C. Slides were incubated with Phospho-p38 for one hour, then with HRP-tagged secondary antibody, DISC. OmniMap Anti-Rb HRP RUO (Roche, 760-4311) followed by ChromoMap DAB detection (Roche, 760-159) on a Roche Ventana Discovery ULTRA. Nuclei were counterstained with Hematoxylin (Roche, 760-2021) for 8 mins and coverslipped. Staining for CTNNB1/β-Catenin (Agilent [Dako], M3539, 1:250): slides were subjected to deparaffinization and antigen retrieval with CC1 (Roche, 950-500) for 24 mins at 95°C. Slides were incubated with Beta-Catenin for 20 minutes, stained with OptiView DAB IHC Detection Kit (Roche, 760-700) on a Roche Ventana Benchmark. Nuclei were counterstained with Hematoxylin (Roche, 760-2021) for 8 mins and coverslipped. All H&E and IHC images were scanned on a Leica AT2 scanner at 40x at UPMC TOPS.

### CD8, CTNNB1, and phospho-p38 IHC quantification

IHC WSI tissue microarray core sections were de-arrayed, cells were segmented and DAB+ cells were detected using QuPath(57) (v0.4.4) (cell segmentation using medianRadiusMircons=1, sigmaMicrons=1.5, Hematoxylin threshold = 0.05; positive cell detection using DAB+ threshold = 0.2 nuclear DAB mean). Tumor regions were annotated on the H&E section of each tumor. All IHC core section images and cell detection objects were then registered to H&E cores using a rigid transform (wsireg(58), v0.3.5). By aligning these cell detections to H&E tumor area, IHC DAB+ cell density was measured inside and outside tumor region of registered cores as the percent of DAB+ / all cells detected for each corresponding IHC core section.

### Multispectral immunofluorescence staining and imaging

Multispectral immunofluorescence staining was performed on 4um tumor FFPE sections: CD3 (Cell Signaling, 85061S), CD8 (Biocare Medical, ACI3160A), PD1 (Abcam, ab137132), PanCK (Santa Cruz Biotech, sc-81714), phospho-p38 (Cell Signaling, 4511S), and DAPI at UPMC TOPS. Automated staining of tissues was performed on the Leica Bond RX. For staining, Akoya Bioscience’s Opal 6-Plex Manual Detection Kit was used according to the manufacturer’s instructions (NEL861001KT). Imaging was performed at 20x on the PhenoImager^TM^ HT (Akoya Biosciences).

### Multispectral immunofluorescence image analysis

Cores were de-arrayed and spectrally unmixed using InForm^®^ (v2.4.6) and Phenochart™ (v1.0) (Akoya Biosciences). Multi-channel composite TIFF files were exported for further data analysis. Cell segmentation was performed on DAPI channel using Stardist(59). Marker positive/negative cells were classified using a machine learning approach. Briefly, a small number of positive/negative cells (30∼50 per class per ROI) were manually selected as the training set to build a Random Forests model, which was then used to predict all remaining cells. Measurement matrices consisting of centroid position (x,y), per-channel intensity, and class label of phenotyped cells were exported and further processed in R (v4.1.2).

### Digital Spatial Profiling (DSP) GeoMX Sample preparation

DSP GeoMx platform (Bruker Spatial Biology; formerly NanoString Inc.) was used for spatial transcriptomics and conducted in the UPMC Hillman Cancer Center Flow Cytometry Facility (FC), same as our previous work(60). 32 HNSCC and 56 LUSC tumors were deparaffinized and stained with GeoMx Solid Tumor TME Morphology Kit including PanCK [Alexa 532, green] and a nuclear stain [Syto-13, blue]. Slides were incubated with The Human Whole Transcriptome Atlas library (WTA) prior to collection. For collection, the GeoMx platform was utilized to select ROIs on the slides. Tumor cell segmentation mask was introduced within each ROI using morphology marker PanCK^+^. Plates were collected and sent for sequencing at the UPMC Children’s Sequencing Facility using standard protocols.

### DSP analysis

Data normalization, quality control (QC), and analysis were conducted by the UPMC Hillman Cancer Bioinformatics Services (CBS)(60), using the methods described in our previous work(60). Sequencing files were converted from FastQ to Digital Count Conversion (DCC) format using The GeoMx NGS Pipeline (v2.0.21). Reads were processed to clip adaptors, merge overlapping mates, align to the Readout Tag Sequence-ID (RTS-ID) barcodes, and remove PCR duplicates by Unique Molecular Identifiers (UMI). The resulting read count matrix was passed through a robust QC procedure to remove ROIs of low surface area (<3000 µm^2^) or low perfect aligned reads (<80%), and probes of low quality or identified as global or local outliers (fails Grubbs outlier test in ≥ 20% segments). The remaining data were upper quartile normalized and log2-transformed. Genes of low expression (<3.5 in HNSCC DSP or <4.5 in LUSC DSP dataset; the threshold is cohort-specific because it was determined by the distribution of negative probe expression [NegProbe-WTX] within each dataset) were removed prior to statistical comparisons. Genes differentially expressed between groups of interest were detected using linear mixed-effects models (LMM) in R (lme4, v1.1-27.1), with group (e.g., high/low phospho-p38 tumors) as the fixed effect and slide id as the random effect. P-values were adjusted using BH-FDR procedure for multiple comparisons.

### Pathway enrichment analysis

Pathways enriched in significant DEGs from HNSCC DSP (FDR 0.10, FC≥1.5 or ≤-1.5) were identified using Enrichr(61, 62) (June 8, 2023) with MSigDB(63) hallmark 2020 gene sets. Enrichment was evaluated by two-sided Fisher’s exact test with multiple comparison adjustment by BH-FDR.

### STRING network analysis

Protein-protein networks were constructed on genes/proteins of interest using STRING(42) functional protein association networks (v12.0), with analysis parameters as: confidence score >0.4; active interaction sources as “Experiments”, “Databases”, and “Co-expression”. Markov Cluster Algorithm(64) (MCL; November 3, 2024) was used to cluster the 23 genes consistently downregulated in high *versus* low phospho-p38 tumor cells from HNSCC and LUSC DSP analysis.

### Human cancer cell Lines and culture condition

HPV-negative human head and neck squamous cell carcinoma (HNSCC) cell lines, PCI 13 and PCI4B were a generous gift from Dr. Robert Ferris. Human lung squamous cell carcinoma (LUSC) cell lines, H520 and H1703 were a generous gift from Dr. Laura A. Stabile, (UPMC Hillman Cancer Center, University of Pittsburgh, PA, USA). PCI13 and PCI4B cells were maintained in IMDM medium supplemented with 10% fetal bovine serum, 1% L-glutamine and 1% NEAA. H520 and H1703 cells were maintained in RPMI with 10% FBS. PCI13, PCI4B, H520, and H1703 cell lines were last authenticated in January of 2023. All cell lines were found to be free from mycoplasma and authenticated stocks of cell lines were used for this study. Empty vector (EV) transfected control cells or human *MAPK14* (p38α) shRNA knockdown cells were generated by stable transfection with EV control (GIPZ lentiviral empty vector shRNA control) or with three different clones of *MAPK14* GIPZ lentiviral shRNA (clones Id; V2LHS_113220; V2LHS_113215; V3LHS_316966) lentiviral plasmids, respectively, and selected by puromycin (0.5 to 2.0 µg/mL) antibiotic. Glycerol stocks of GIPZ Lentiviral EV shRNA control (Catalog No: RHS4349) and three different GIPZ Human *MAPK14* shRNA plasmids (1-Catalog No: RHS4430-200212538: Clone Id: V2LHS_113215; 2-Catalog No: RHS4430-200183464; Clone Id: V2LHS_113220; 3 - Catalog No: RHS4430-200266677: Clone ID: V3LHS 316966) were purchased from Horizon Discovery Group plc.

### Western Blot

Cells were washed twice with ice-cold PBS, lysed on ice in cell lysis buffer (RIPA buffer). The cell lysate were cleared by centrifugation at 14 000 g for 20 minutes and protein content was quantified using Bradford method and equal amount of protein was subjected to sodium dodecyl sulfate-polyacrylamide gel electrophoresis. Proteins were transferred onto polyvinylidene fluoride membrane. After blocking with 5% non-fat dry milk in Tris-buffered saline containing 0.05% Tween-20, the membrane was incubated with the p38 (1:20,000 dilution; Cell Signaling Technology: catalog number # 9212) or phospho-p38 (1:1000 dilution; Cell Signaling Technology: catalog number # 4511) primary antibody for overnight at 4^0^C. Subsequently, the membrane was incubated with the appropriate secondary antibody, and the immunoreactive protein bands were visualized using chemiluminescence method. Each membrane was stripped and re-probed with housekeeping protein β-Actin (1:100,000 dilution; Sigma Aldrich: catalog number # A5441) to correct for differences in protein loading. Change in protein level was quantified using densitometric scanning of the immunoreactive band and corrected for β-Actin loading control by UN-SCAN-IT software (Silk Scientific).

### Pharmacologic p38 inhibition with pexmetinib

Pexmetinib (ARRY-614) was obtained from Pfizer, Inc. While some work describes it as a p38 /Tie2 inhibitor, pharmacologic studies (Array Biopharma, acquired by Pfizer) demonstrated that pexmetinib has 10-fold greater potency against p38 MAPK relative to Tie2. In human whole blood assays in vitro, the predicted plasma concentrations to achieve 50% target inhibition (IC_50_), after protein binding correction, were 172 nM for phospho-p38 compared with 2282 nM for phospho-Tie2. Consistent with this, in vivo pharmacodynamic studies in murine models showed that p38 inhibition was achieved at substantially lower concentrations than those required for Tie2 inhibition (IC50 = 203 nM and 2066 nM, respectively) (30, 31). Based on these data, pexmetinib is not considered a clinically relevant Tie2 inhibitor at pharmacologically achievable exposures. Instead, it functions as a highly selective p38 MAPK inhibitor and was therefore used in this study as a pharmacologic agent to validate p38 pathway inhibition in human cancer cell lines.

### Preparation of tumor conditioned media

5-8 x 10^5^ human cells were plated in triplicate in 60 mm dishes. After overnight incubation, the media were replaced with either DMSO or pexmetinib (p38 inhibitor:10 nM), and cells were incubated for an additional 24 hours. Following this incubation, media were collected and centrifuged at 3500 x g for seven minutes, and the supernatant was collected. The supernatant was further centrifuged using a 3 kDa Amicon® Ultra Centrifugal Filter (Millipore Sigma, catalog number # UFC800324) at 4000 rpm for 10 minutes. After centrifugation, the supernatants were collected as conditioned media and stored at-80°C for downstream experiments. Separately, EV control and p38α shRNA-stably transfected human cancer cells were plated in triplicate in 60 mm dishes and incubated for 48 hours. After incubation, media were collected, centrifuged at 3500 x g for seven minutes, and the supernatant was collected and stored at-80°C.

### CD8^+^ T cell migration assay

Human CD8^+^ T cells were isolated from peripheral blood mononuclear cells (PBMCs) obtained from healthy donor leukopacks (Vitalant, Scottsdale, AZ), using CD8^+^ T (Catalog number: 130-096-495; Miltenyi Biotec) cell isolation kits following the manufacturer’s instructions. The purity of the isolated CD8^+^ T cells was confirmed by flow cytometry using live/dead, CD45^+^, CD3^+^, and CD8^+^ staining markers. A total of 5 × 10^5^ CD8^+^ T cells were then resuspended in 100 µL of T migration medium (RPMI supplemented with 1% FBS, 10 mM HEPES buffer, pH 6.9) and placed in the upper chamber of a 96-well transwell plate with a 5-µm pore polycarbonate membrane (Corning). The lower chamber contained conditioned medium from EV control cells, p38α shRNA-transfected cells, or serum-free media (negative control). T cells were allowed to migrate to the lower chamber over 18 hours at 37°C in a CO₂ incubator, and migrated cells were counted manually using trypan blue and a hemocytometer. To ensure adequate migrated-cell numbers while maintaining robust assay reproducibility, we selected an 18-hour incubation period for conditioned-media Transwell migration assays, which falls within the range commonly used for this type of experiments in literature(65, 66).

For migration assays with DMSO-or pexmetinib-treated human cancer cell lines (HNSCC cell lines PCI4B and PCI13, LUSC cell lines H1703 and H520) conditioned media, isolated T cells were pre-cultured in RPMI 1640 medium containing 10% FBS, 1% L-glutamine, 1% penicillin-streptomycin, 1% HEPES, IL-2 (30 IU/mL), and β-mercaptoethanol (50 µM) for 72 hours to stimulate T cell activation. Conditioned media from treated cell lines were used in the lower chambers. Additionally, CXCL16 neutralizing antibody (30 ng/mL, catalog #AF976) and control goat IgG (30 ng/mL, catalog #AB-108-C) were included in the conditioned media in specific experimental groups, and pertussis toxin (200 ng/mL, Gibco catalog #MAN0004270) was used as a negative control. The transmigration assay was conducted for 18 hours, same as above.

### Cell harvesting for RNAseq experiments

The EV control and shRNA KD cells from human HNSCC (PCI4B and PCI13), human LUSC (H1703 and H520) were harvested for RNAseq. Cells were plated in 6 cm dishes at a density of 4 × 10^5^ to 8 × 10^5^ cells/dish in triplicate and were incubated for 24hours. After incubation cells were harvested by trypsin treatment and total RNA was extracted using the RNeasy mini kit (Qiagen, Germantown, MD; Cat. No. 74104). RNA quality was determined by RNA integrity number and Qubit^®^.

### RNAseq library preparation, sequencing, and data analysis

RNAseq library preparation and sequencing were performed at Novogene Inc. on an Illumina NovaSeq instrument. Paired-end (PE) 150bp reads were generated, with an average of 20 million read pairs per sample. Data was analyzed following established protocols(67). After quality control (QC), PE reads were pseudoaligned against human reference transcriptome (GRCh38 assembly, Gencode v38 annotation) using Kallisto (v0.46.2), summarized at gene level using tximport (v1.20.0), followed by TMM normalization and log_2_ transformation. Lowly expressed genes (counts per million of mapped reads [CPM]<3) were excluded prior to statistical comparisons. Differentially expressed genes (DEGs) between groups of interest were detected using limma voom with precision weights(68) (v3.54.2), with p-values adjusted by BH-FDR method.

### RNA isolation and real-time reverse transcription polymerase chain reaction (RT-PCR)

The expression of *MAPK14* and *CXCL16* was determined by RT-PCR. Total RNA from EV control and shRNA KD cells or DMSO or pexmetinib treated cells was isolated using RNeasy kit. One µg RNA was used for cDNA synthesis with the use of Applied Biosystems™ High-Capacity cDNA Reverse Transcription Kit (Catalog number # 43-688-14). Quantitative PCR was performed using Bullseye Real Time qPCR Master Mix (EvaGreen) for 40 cycles. The expression of *MAPK14* and *CXCL16* were normalized to glyceraldehyde 3-phosphate dehydrogenase (*GAPDH*). The primers for human *MAPK14, CXCL16,* and *GAPDH* were as follows: Forward (MAPK14): 5’-CAGTGGGATGCATAATGGCC-3’; reverse (*MAPK14*) 5’-GCATCTTCTCCAGCAAGTCG-3’; forward (*CXCL16*): 5’-AGCTCACTCGTCCCAATGAA-3’; reverse (*CXCL16*): 5’-ACAGCACATAGGAAAGGGCT-3’; 5’-forward (*GAPDH*): 5’-GGACCTGACCTGCCGTCTAGAA-3’; reverse (*GAPDH*): 5’-GGTGTCGCTGTTGAAGTCAGAG-3’. The PCR conditions were as follows: 95°C for 10 minutes followed by 40 cycles of 95°C for 15 seconds, 60°C (*MAPK14* and *GAPDH*) and 58°C (*CXCL16*) for 1 minute, and 72°C for 30 seconds.

### Quantitation of CXCL16 in cell culture medium

Conditioned media from DMSO-treated control and pexmetinib (10 nM)-treated human cell lines (PCI13, PCI4B, H520, and H1703), as well as from stably transfected EV control and corresponding p38 shRNA cell lines (PCI13, PCI4B, H520, and H1703), were used to assess CXCL16 protein release. Equal volumes of conditioned media were analyzed for CXCL16 levels using the Human CXCL16 ELISA Kit from RayBiotech (catalog #ELH-CXCL16), following the manufacturer’s instructions.

### Statistical analysis

Measurements were taken from distinct samples. The Wilcoxon signed-rank test was used to compare expression values between tumor groups. Bulk RNAseq DEG analysis between groups was performed using empirical Bayes regression models in limma voom with precision weights. scRNAseq tumor cell expressing pathway score comparisons between tumor groups were performed using linear mixed-effects models in lme4, with tumor id as random effect and group of interest as fixed effect (formula: `pathway_score ∼ 0 + tumor_group + (1 | tumor_name), RMEL=FALSE`). scRNAseq pathway score comparisons between malignant epithelial cells and fibroblasts were performed using linear mixed-effects models with a nested design (formula: `pathway_score ∼ 0 + cell_type + (1 | tumor_name/cell_type), RMEL=FALSE`). REML was set to FALSE to use maximum likelihood for model fitting, and likelihood ratio test (LRT) was used to compute p-values. For DEG detection from DSP data, linear mixed-effects model was used with tumor group as the fixed effect and slide id as the random effect (formula: `dsp_gene_expression_norm ∼ 0 + tumor_group + (1 | slide_id)`). In addition, statistics for specific analysis were described above or in figure legends. FDR is controlled at 0.10 by the BH FDR procedure(69) unless otherwise noted. All tests are two-sided. Statistical analysis was performed using R (v4.1.2) and Bioconductor (release 3.14).

## LIST OF ABBREVIATIONS

CPTAC: Clinical Proteomic Tumor Analysis Consortium
CTNNB1: Beta-catenin
DC: dendritic cell
DSP: digital spatial profiling
FDA: Food and Drug Administration
HPV: human papillomavirus
HNSCC: Head and Neck Squamous cell carcinoma
ICGC: International Cancer Genome Consortium
ICI: immune-checkpoint inhibitor
IFN: interferon
IHC: immunohistochemistry
LUAD: lung adenocarcinoma
LUSC: lung squamous cell carcinoma
MAPK: mitogen-activated protein kinase
mIF: multispectral immunofluorescence
PD-L1: programmed death-ligand 1
PD-1: programmed cell death protein 1
RCC: renal clear cell carcinoma
RNAseq: Ribonucleic acid sequencing
scRNA-seq: single cell ribonucleic acid sequencing
SKCM: Skin Cutaneous Melanoma
TCGA: The Cancer Genome Atlas
TMA: Tumor microarrays
TMB: tumor mutational burden
TME: Tumor microenvironment
WNT: Wingless-related integration site

## DECLARATIONS

### Ethics approval and consent to participate

Human specimens were obtained from patients consented under The University of Pittsburgh institutional review board (IRB)-approved protocol (HCC 99-069 for head and neck, and STUDY12070229 for lung). Written informed consent was obtained from all participants before the study.

### Consent for publication

All authors have read and approved the final manuscript for publication.

### Data availability

The deidentified clinical data and processed biological data are provided in supplementary tables. This includes the cell densities from PhenoImager^TM^ HT experiments, and DEG statistics from DSP GeoMX experiments. Sequencing data generated in this study will be deposited to NCBI GEO repositories. Public available data were cited with accession numbers for the datasets as appropriate.

### Code availability

This paper utilizes open-source software/tools and libraries as described in Methods. Any additional information required to reanalyze the data reported in this paper is available from the corresponding authors upon request.

## Funding

This work was supported by National Institutes of Health (NIH) grant R01DE031729 (R.B., D.P.Z.), DoD-MRP TSA ME240329 (R.B.), P50CA097190 (R.B., R.L.F., H.D.S., D.P.Z.), NIH P50CA254865 (R.B.), NIH P50CA272218 (R.B.), R01DE028061 (H.D.S.), RO1DE028105 (H.D.S.), RO1DE032337 (H.D.S.), P50CA254865 (R.B., D.D.), American Lung Association Innovator Award IA-1273956 (R.B.), LCRF Leading Edge Award (R.B.), Cancer Immunology Training Program T32CA082084 (R.E.D.), the University of Pittsburgh Liver Research Center (PLRC award to R.E.D, part of NIH P30 DK120531), University of Pittsburgh Cancer Cell Biology Program (CBP from NIH CCSG P30CA047904 to R.E.D.) and in part by National Cancer Institute (NCI) through the UPMC Hillman Cancer Center (HCC) CCSG award P30CA047904 (R.L.F., R.B.), and The University of Pittsburgh CRCD through the resources provided, specifically the HTC clusters supported by NIH S10OD028483. Pfizer Inc. provided drug substance but had no input on the study design or the interpretation of the results.

### Authors’ contributions

R.B. conceived the study and supervised the project. RB acquired the data, developed the methodology, and analyzed the data. R.E.D. revised methods and performed analysis. KBS performed the in vitro experiments and data analysis. R.D. performed the cell viability assay from p38i in cell lines. R.S. and C.K. maintained the cell lines, and assisted with data analysis from experiments. B.I. and C.D. assisted with the spatial analysis. S.N. managed the project and acquired specimens. M.J. performed the multispectral IF staining in human specimens. K.S. performed sample processing, H&E staining, IHC staining, and image scanning. C.R. and A.K. assisted with clinical data. A.P.S., E.M.M., and T.C.B. performed the DSP experiment. Q.G., C.Z., A.D.S., and R.R.S. performed the pathology annotation and review. A.C.S., R.D., and L.V. assisted with the design of immune-related experiments. L.V. contributed critical feedback on data interpretation. H.D.S. facilitated specimen inquiries. L.P.S. contributed the lung specimens. R.L.F. and D.P.Z. contributed the head and neck specimens and data from UPitt ICI cohort. R.E.D., R.B., and J.J.L. interpreted the results. RB and JJL wrote the manuscript. All authors contributed to the final manuscript. All authors read and approved the final manuscript.

## Acknowledgements.

We thank the patients and families for their participation in this study. We thank Dr. Fangping Mu for technical assistance at The University of Pittsburgh Center for Research Computing and Data (CRCD) high-performance computing clusters (HPC), and Shaila Fye for the initial assessment of *Puram et al.* scRNAseq data. This project used the UPMC HCC Cancer Bioinformatics Facility (CBS), Translational Oncologic Pathology Services (TOPS), and Flow Cytometry Facility (FC).

