## Supplementary material for "Integrative multi-omic analysis identifies tumor-intrinsic p38 as a driver of immune exclusion in human epithelial cancers": Supp figs

This document contains **Supplementary Figures 1 to 6** and **Title of Supplementary Tables**. The **Supplementary Tables 1 to 11** were supplied as separate spreadsheets**.**

### Supplementary Figures

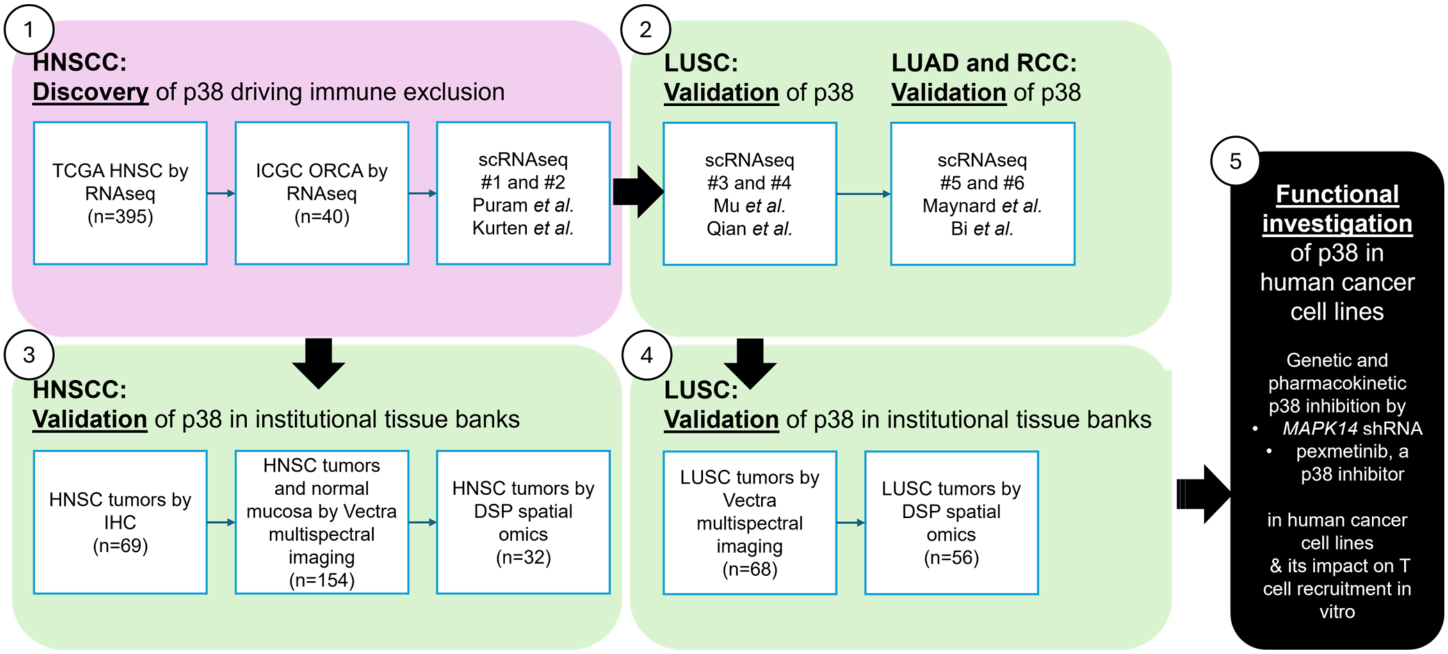


**Supplementary Figure 1. Overall schema illustrating the discovery/validation workflow of the study design.**

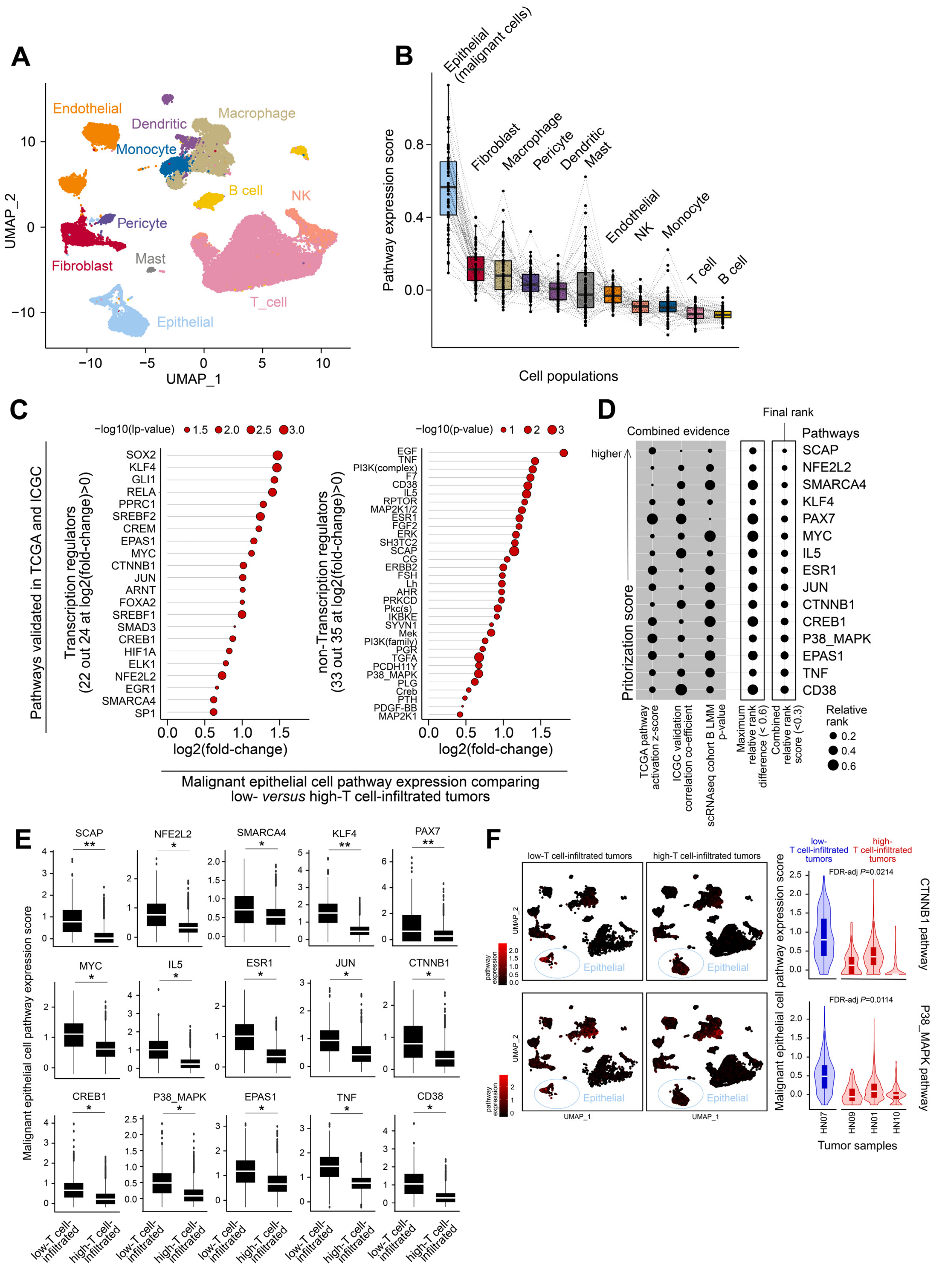
A
B
C
D
E
F
Endothelial
Dendritic
Macrophage
Monocyte
B cell
NK
Pericyte
Fibroblast
Mast
T_cell
Epithelial
UMAP_1
UMAP_2
Epithelial (malignant cells)
Fibroblast
Macrophage
Pericyte
Dendritic
Mast
Endothelial
NK
Monocyte
T cell
B cell
Pathway expression score
Cell populations
Pathways validated in TCGA and ICGC
Transcription regulators (22 out 24 at log2(fold-change)>0)
non-Transcription regulators (33 out 35 at log2(fold-change)>0)
−log10(lp-value)
−log10(p-value)
log2(fold-change)
Malignant epithelial cell pathway expression comparing low- versus high-T cell-infiltrated tumors
Combined evidence
Final rank
Pathways
higher
Pritorization score
TCGA pathway activation z-score
ICGC validation correlation co-efficient
scRNAseq cohort B LMM p-value
Maximum relative rank difference (< 0.6)
Combined relative rank score (<0.3)
Relative rank
SCAP
NFE2L2
SMARCA4
KLF4
PAX7
MYC
IL5
ESR1
JUN
CTNNB1
CREB1
P38_MAPK
EPAS1
TNF
CD38
Malignant epithelial cell pathway expression score
low-T cell-infiltrated
high-T cell-infiltrated
low-T cell-infiltrated tumors
high-T cell-infiltrated tumors
pathway expression
low- T cell-infiltrated tumors
high- T cell-infiltrated tumors
FDR-adj P=0.0214
FDR-adj P=0.0114
CTNNB1 pathway
P38_MAPK pathway
HN07
HN09
HN01
HN10
Tumor samples

**Supplementary Figure 2. non-T cell-inflamed phenotype-associated pathways are activated in malignant cells from HPV-negative HNSCC of low T cell infiltration from scRNAseq cohort #2 (Kurten et al.).** (**A**) Distribution of tumor, stroma, and immune cell subsets on UMAP (n=41,435 cells from nine tumors). (**B**) Expression of 59 pathways from **Fig. 1D** across cell populations. Pathway scores were computed as the average expression of all genes involved in a pathway. Each data dot is the average score of a pathway in each cell population. Dotted lines connect the per-cell population scores of the same pathway. (**C**) 55 out of 59 pathways showed higher expression in 689 malignant epithelial cells of low-T cell-infiltrated relative to 2,834 malignant epithelial cells of high-T cell-infiltrated tumors. (**D**) 15 pathways that passed prioritization score (combined relative rank) < 0.6 and maximum relative rank difference < 0.3. Six out of 15 pathways overlap with the top pathways from the Puram cohort in **Fig. 2D**. For each of the 59 pathways from **Fig. 1D**, its combined relative rank was computed as the geometric mean of three values: the relative rank in TCGA pathway activation (z-score higher to lower), the relative rank in ICGC anti-correlation with T cell-inflamed expression (coefficient highly to lowly negative), and the relative rank in scRNAseq pathway expression comparing malignant epithelial cells from low-T cell-infiltrated *versus* high-T cell-infiltrated tumors (p-values smaller to larger). (**E**) Expression of the 15 pathways in the same order as shown in **D**. (**F**) Expression of CTNNB1 pathway and p38 pathway from low- *versus* high-T cell-infiltrated tumors. Four out of nine tumors with at least 40 malignant epithelial cells per sample were included in analysis. Linear mixed-effects model via maximum likelihood was used in **C**, **E** and **F**, with tumor group as the fixed effect and tumor id as the random effect. Likelihood ratio test (LRT) was used with the fitted model for computing p-values, followed by BH-FDR correction for multiple comparisons. Denotation: ** FDR-adjusted $P<0.01$, * FDR-adjusted $P<0.05$, otherwise the numbers are shown. For boxplots in **B**, **E**, and **F**, the median is shown by the horizontal center line, with the 25th and 75th percentiles depicted by the boxes; the whiskers extend to the outermost data point within a range of up to 1.5 times the interquartile range (IQR).

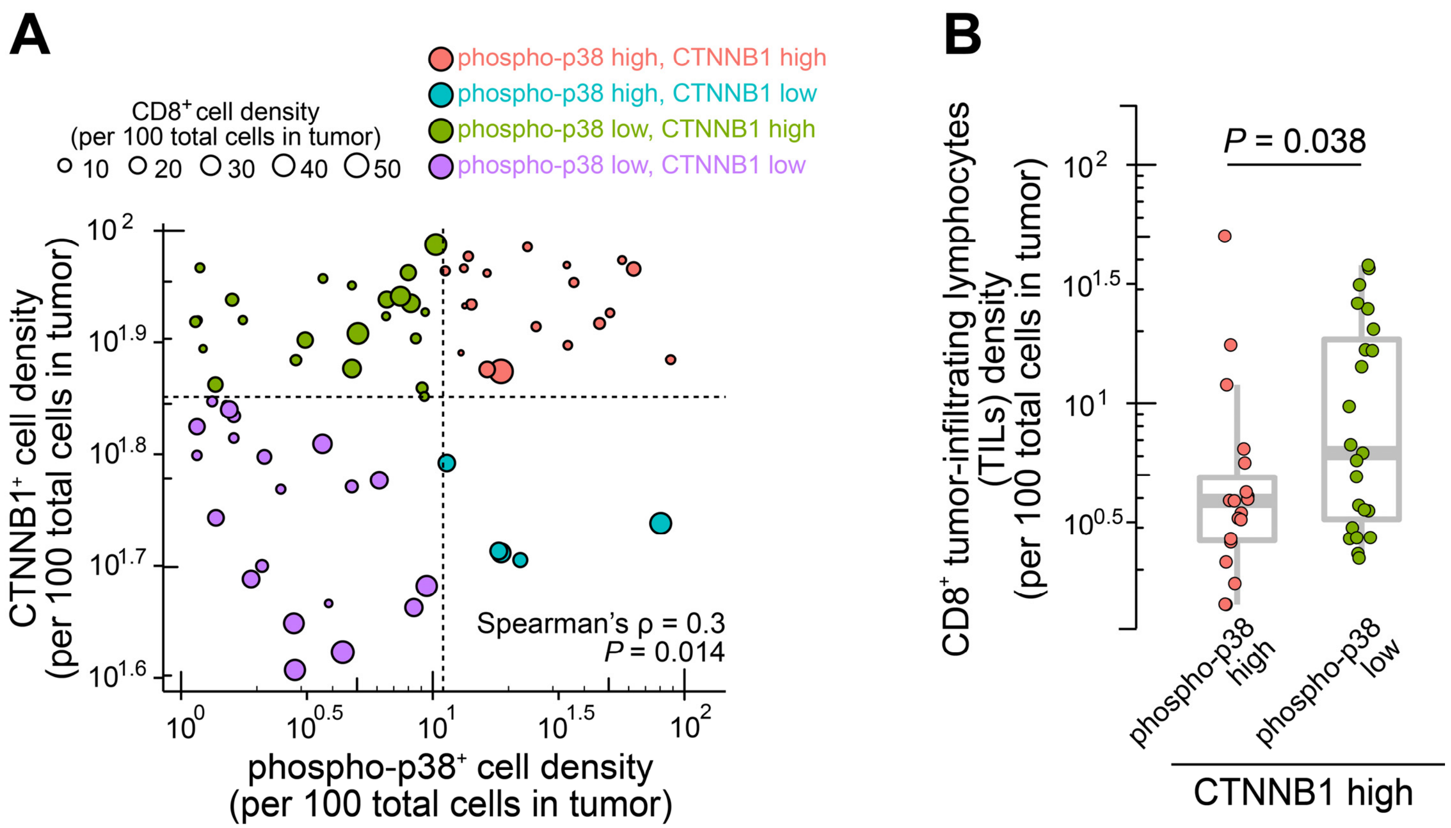


**Supplementary Figure 3**. **IHC quantification of CTNNB1$^+$, phospho-p38$^+$, or CD8$^+$ cell density in HNSCC tumor.** (**A**) Correlation between phospho-p38$^+$ cell density and CTNNB1$^+$ cell density. n=69 HNSCC cores are shown. Each data point represents one core, with color representing the four categories split by CTNNB1$^+$ cell density high/low or phospho-p38$^+$ cell density high/low, and size representing the CD8$^+$ cell density. (**B**) Comparison of CD8$^+$ cell density between phospho-p38$^+$ high and low groups in the context of CTNNB1-high tumor cores from **A**. The phospho-p38$^+$ high group was defined as having ≥10 phospho-p38$^+$ cells per 100 total cells, and CTNNB1-high tumor cores were defined as having ≥70 CTNNB1$^+$ cells per 100 total cells. Spearman's correlation was used in **A**, two-sided Wilcoxon rank sum test in **B**.

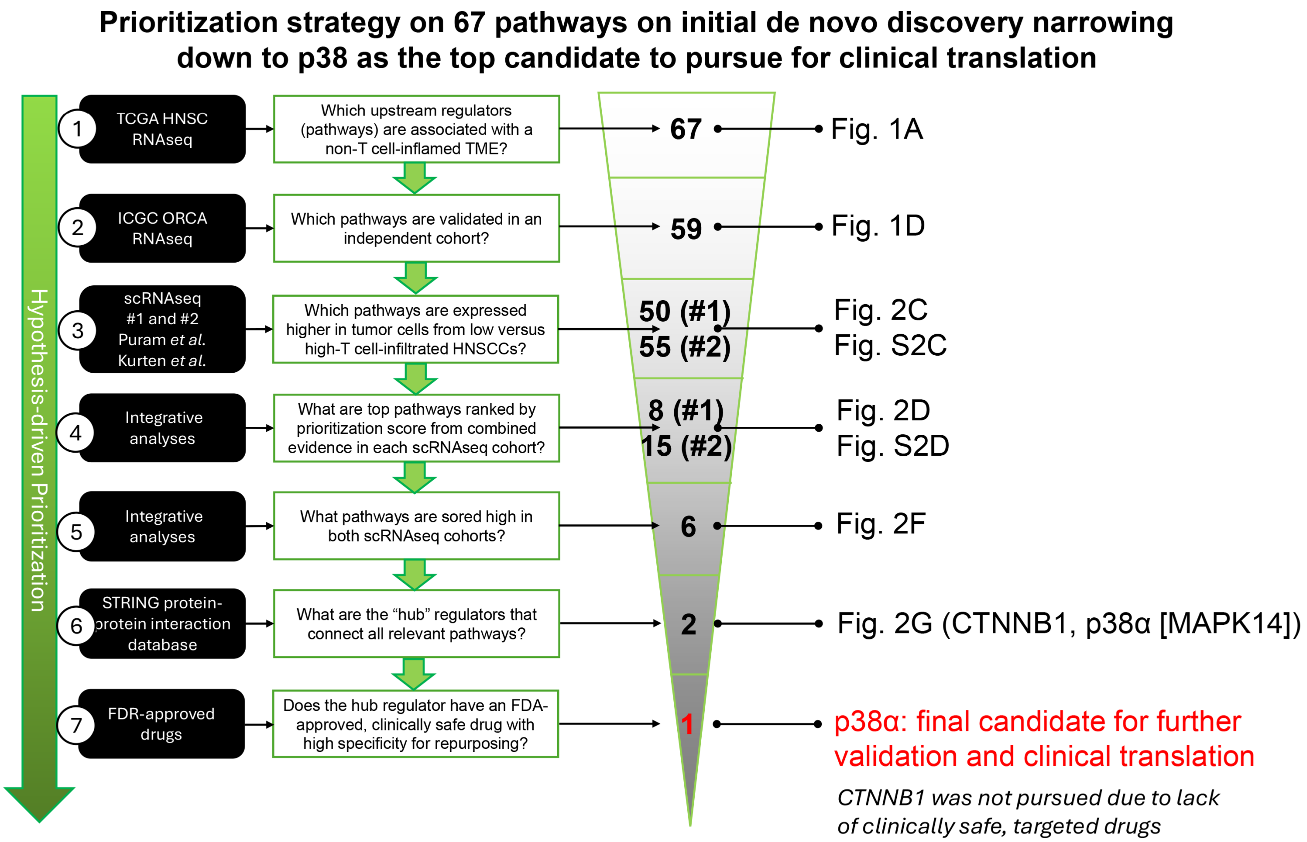


**Supplementary Figure 4. Prioritization strategy starting with a discovery of 67 pathways narrowing down to p38 as the top candidate for clinical translation.**

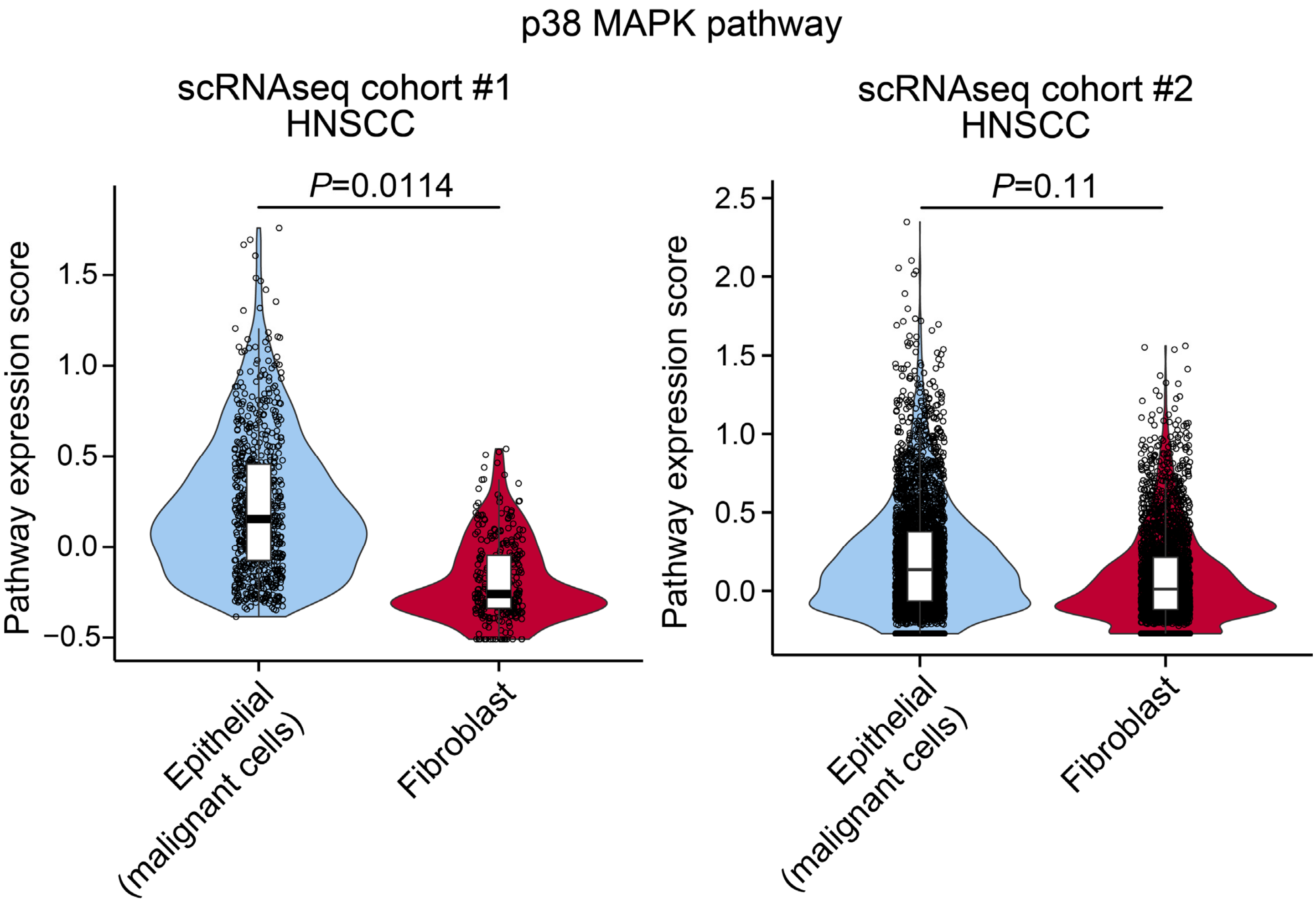


**Supplementary Figure 5. p38 pathway expression scores in malignant epithelial cells and fibroblasts from both HPV-negative HNSCC scRNAseq cohorts.** P-value was computed by linear mixed-effects models (LMM) with a nested design (tumor id as the blocking factor), followed by likelihood ratio test (LRT). For boxplot, the median is shown by the horizontal center line, with the 25th and 75th percentiles depicted by the boxes; the whiskers extend to the outermost data point within a range of up to 1.5 times the interquartile range (IQR).

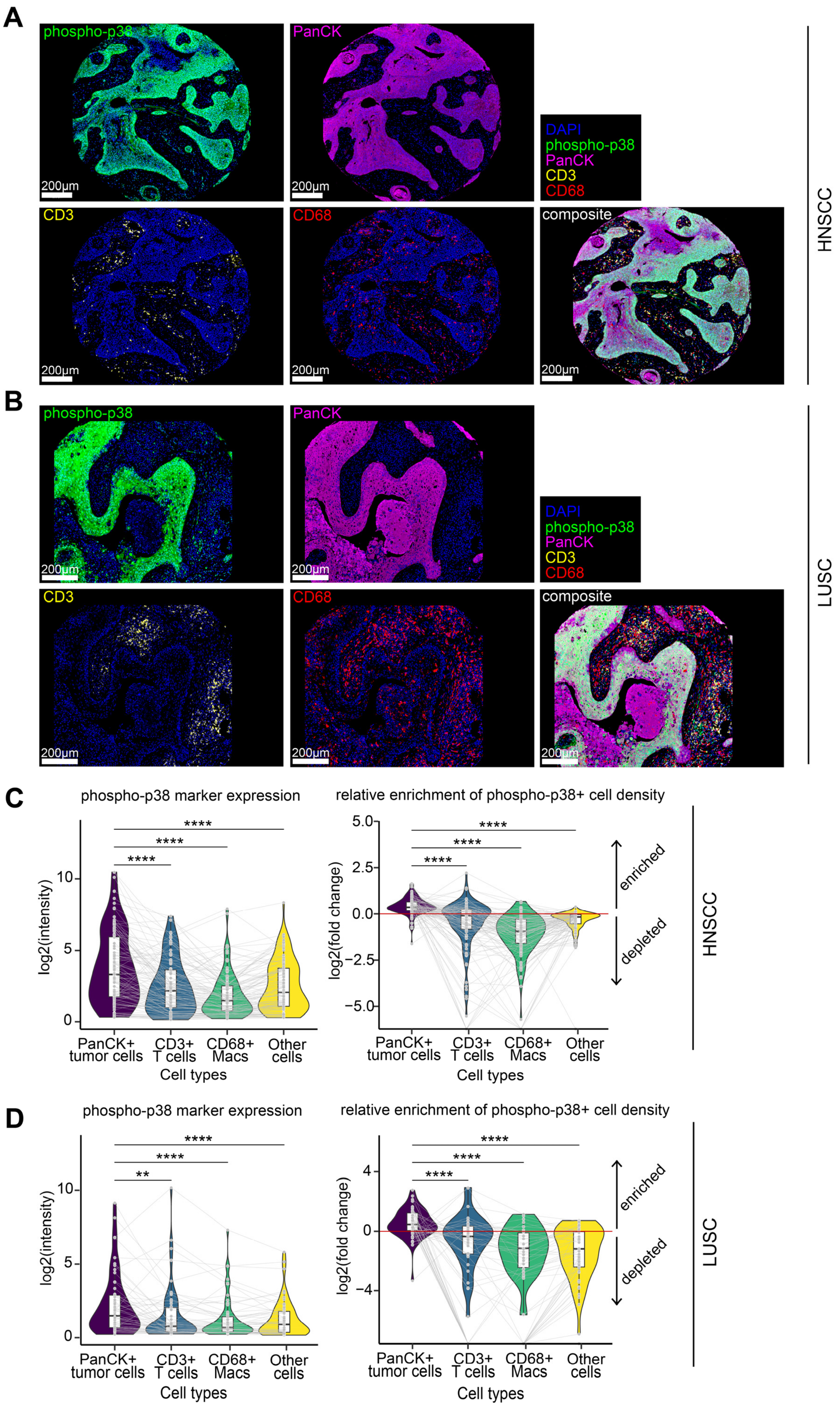
A
phospho-p38
PanCK
CD3
CD68
composite
DAPI
phospho-p38
PanCK
CD3
CD68
200µm
HNSCC
B
LUSC
C
phospho-p38 marker expression
relative enrichment of phospho-p38+ cell density
log2(intensity)
log2(fold change)
enriched
depleted
PanCK+ tumor cells
CD3+ T cells
CD68+ Macs
Other cells
Cell types
****
**
D

**Supplementary Figure 6. Phospho-p38 expression is dominantly detected in tumor cells from HNSCC and LUSC human tumor samples by multispectral immunofluorescence Vectra Polaris.** (**A-B**) Representative examples of phospho-p38, PanCK, CD3, and CD68 staining of HNSCC in **A** and LUSC in **B**. (**C-D**) Phospho-p38 in HNSCC in **C** and LUSC in **D**. (**left**) Distribution of phospho-p38 marker expression (imaging intensity) across major cell types in tumor. Each dot represents one tumor sample. For each sample, the average of $\log_2$-transformed phospho-p38 single-cell intensity values from that sample is shown on *y*-axis. *x*-axis represents main cell types: $PanCK^+$ tumor cells, $CD3^+$ T cells, $CD68^+$ macrophages, and other cells. Gray lines connect data from the same sample. $PanCK^+$ tumor cells exhibited significantly higher phospho-p38 expression compared to immune and other populations in both HNSCC and LUSC tumors. (**right**) Relative enrichment of phospho-$p38^+$ cells across cell types. For each sample, we calculated the observed fraction of phospho-$p38^+$ cells belonging to each cell type, and compared this to the expected fraction based on that cell type's overall abundance. Enrichment was defined as $\log_2$(fold change of [observed/expected]) for each cell type per sample. A value > 0 indicates overrepresentation (enrichment), and < 0 indicates underrepresentation (depletion). Tumor ($PanCK^+$) cells were significantly enriched for phospho-$p38^+$ cells, whereas immune and other cells were depleted for phospho-$p38^+$ cells. Together, these results highlight tumor epithelial cells as the dominant contributors to phospho-p38 activation in HNSCC and LUSC. P-value by two-sided Wilcoxon signed-rank test in **C** and **D**. BH-FDR adjusted p-values are shown. Denotation: **** $P<0.0001$, *** $P<0.001$, ** $P<0.01$, * $P<0.05$.

**Supplementary Tables**

**Supplementary Table 1**. List of drugs targeting the 59 oncogenic pathways associated with immune exclusion in HPV-negative HNSCC identified in our study from The Drug Gene Interaction Database (DGIdb).

**Supplementary Table 2.** 59 pathways in HNSCC scRNAseq cohort A comparing low- vs high-T cell-infiltrated tumors.

**Supplementary Table 3**. 59 pathways in HNSCC scRNAseq cohort B comparing low- vs high-T cell-infiltrated tumors.

**Supplementary Table 4.** Pathologist-evaluated lymphocyte-based inflammation scores from H&E images of cohort #2 (Kurten et al.).

**Supplementary Table 5.** Ensemble scores of 59 pathways from TCGA, ICGC and both HNSCC scRNAseq cohorts.

**Supplementary Table 6.** IHC measures of CD8+, CTNNB1+, and phospho-p38+ cell density from HNSCC tissues.

**Supplementary Table 7.** Multispectral imaging measures of CD3+, CD8+, PanCK+, and phospho-p38+ cell density from HNSCC tissues.

**Supplementary Table 8**. DEGs from HNSCC tumor DSP comparing phospho-p38 high versus low tumor cells (fold change ≤ -1.5 or ≥ 1.5, FDR-adjusted $P<0.10$).

**Supplementary Table 9**. Pathway analysis of the HNSCC tumor DSP DEGs using Enrichr with MSigDB hallmark genesets.

**Supplementary Table 10**. Multispectral imaging measures of CD3+, CD8+, PanCK+, and phospho-p38+ cell density from LUSC tissues.

**Supplementary Table 11**. 23 genes consistently downregulated in phospho-p38 high PanCK+ ROIs in both HNSCC tumor DSP and LUSC tumor DSP datasets.
